# A Genomic Perspective on the Evolutionary Diversification of Turtles

**DOI:** 10.1101/2021.10.14.464421

**Authors:** Simone M. Gable, Michael I. Byars, Robert Literman, Marc Tollis

**Affiliations:** School of Informatics, Computing, and Cyber Systems, Northern Arizona University, PO Box 5693, Flagstaff, AZ 8601, USA; Department of Biological Sciences, University of Rhode Island, 120 Flagg Road, Kingstown, RI, 0288, USA

**Keywords:** turtles, genomes, phylogeny, discordance

## Abstract

To examine phylogenetic heterogeneity in turtle evolution, we collected thousands of high-confidence single-copy orthologs from 19 genome assemblies representative of extant turtle diversity and estimated a phylogeny with multispecies coalescent and concatenated partitioned methods. We also collected next-generation sequences from 26 turtle species and assembled millions of biallelic markers to reconstruct phylogenies based on annotated regions from the western painted turtle (*Chrysemys picta bellii*) genome (coding regions, introns, untranslated regions, intergenic, and others). We then measured gene tree-species tree discordance, as well as gene and site heterogeneity at each node in the inferred trees, and tested for temporal patterns in phylogenomic conflict across turtle evolution. We found strong and consistent support for all bifurcations in the inferred turtle species phylogenies. However, a number of genes, sites, and genomic features supported alternate relationships between turtle taxa. Our results suggest that gene tree-species tree discordance in these datasets is likely driven by population-level processes such as incomplete lineage sorting. We found very little effect of substitutional saturation on species tree topologies, and no clear phylogenetic patterns in codon usage bias and compositional heterogeneity. There was no correlation between gene and site concordance, node age, and DNA substitution rate across most annotated genomic regions. Our study demonstrates that heterogeneity is to be expected even in well resolved clades such as turtles, and that future phylogenomic studies should aim to sample as much of the genome as possible in order to obtain accurate phylogenies for assessing conservation priorities in turtles.

## Introduction

Next-generation sequencing has greatly increased the numbers of sampled loci for phylogenomic analyses (Lemmon et al. 2012; Faircloth et al. 2013; Edwards et al. 2016), shedding new light on the branching order of diversification across the history of life. However, phylogenetic studies based on genome-scale data often contain a great deal of heterogeneity in the inferred patterns (Hime et al. 2021; Lopes et al. 2021; Morales-Briones et al. 2021; Singhal et al. 2021), particularly in the form of discordance among sites and gene trees (Kumar et al. 2012). Gene tree-species tree discordance occurs when gene trees are in conflict with the underlying species relationships, while site discordance occurs when a subset of sites support different bifurcations in a species tree, and both factors can lead to biased phylogenetic estimates. Understanding patterns and potential sources of phylogenomic heterogeneity will not only settle phylogenetic arguments and reduce bias in tree building, but also provide better models with which to understand trait evolution and help define conservation priorities through comparative methods (Garland et al. 2005; Colston et al. 2021).

Biological factors driving phylogenomic heterogeneity include: differing evolutionary rates across the genome, incomplete lineage sorting, and hidden paralogy (Pamilo and Nei 1988; Galtier and Daubin 2008). Artifacts that drive heterogeneity include sequencing errors and substitution model violations such as genetic saturation, compositional heterogeneity, and codon usage bias (Foster 2004; Cooper 2014; Cox et al. 2014). Accounting for gene and site discordance in phylogenomic estimation often requires partitioning schemes which try to account for differing evolutionary rates and model parameters (Kubatko and Degnan 2007), methods that are consistent with the multispecies coalescent accounting for population-level processes such as incomplete lineage sorting (Edwards et al. 2016), as well as strict filtering of sequence alignments that aim to reduce potential errors. The application of hundreds or even thousands of loci is often an attempt to overcome biases stemming from or driving phylogenomic conflict, and these studies often boast strong branch support as measured by bootstrap replicates or posterior probabilities, likely as a result of reduced sampling variance in large datasets (Minh et al. 2020a). However, competing phylogenomic studies of the same clades that use different datasets and tree building methods can produce conflicting results, despite high levels of statistical support (see modern birds: Jarvis et al. 2014; Prum et al. 2015).

An emerging advantage of the genomic era is the theoretical ability to access the whole genome in order to sample loci representing a diversity of evolutionary rates and coalescent histories (Wolf et al. 2002; Rokas et al. 2003). The most widely used sampling methods include the use of large numbers of coding sequences (Singhal et al. 2021), anchored hybrid enrichment loci (AHE; Lemmon et al. 2012), ultra-conserved elements (UCEs; Faircloth et al. 2013), and transcriptomes (Irisarri et al. 2017), but these are each still reduced representations of a relatively small proportion of the genome (Lynch 2007). Meanwhile, the development of bioinformatics tools and sequencing technologies have enabled chromosome-scale genome assemblies for non-model organisms (see http://dnazoo.org, Dudchenko et al. 2017), potentially providing insights to the genomic regions or types of loci that drive heterogeneous phylogenetic results. Thus, by accessing complete genomes and studying heterogeneity closely, we can more fully comprehend conflicting results across gene trees, obtain more accurate perspectives of species relationships, and gain a fuller understanding of molecular evolution across the genome.

Here, we examined the importance of genome-wide heterogeneity in the phylogenomic estimation of turtles (Order Testudines). Turtles are a near-globally distributed and morphologically distinct clade of shelled reptiles, with a rich fossil history dating from the Triassic Period (Gaffney 1980; Joyce 2007; Joyce et al. 2021). Despite their persistence in the fossil record and across modern ecosystems, almost half of turtle species are listed in the International Union for Conservation of Nature (IUCN) as endangered, critically endangered, or vulnerable to extinction (IUCN, 2021). The approximately 350 extant species of turtles are classified into two main groups, Pleurodira (“side-necked”) and Cryptodira (“hidden-necked”), which are further divided into 8 superfamilies and 14 families (Uetz et al. 2021). Early molecular phylogenetic results disagreed in the placement of some turtle taxa; in particular, the position of the monotypic big-headed turtle (*Platysternon*) and its relationships with or within Testudinoidea (Shaffer et al. 1997; Krenz et al. 2005). Other questions about deeper turtle relationships such as the position of soft-shell turtles (Trionychoidea), the Americhelydiae, and the position of turtles in the amniote phylogeny persisted into the next-generation sequencing era (Chiari et al. 2012; Crawford et al. 2012, 2015; Brown and Thomson 2017). Since then, phylogenomic studies of turtles have produced fully resolved species trees with 100% statistical support at every node and near complete agreement across studies as measured by bootstrap replicates or posterior probabilities (Crawford et al. 2015; Shaffer et al. 2017).

Phylogenomic heterogeneity has been examined in vertebrate groups such as mammals (Tarver et al. 2016; Liu et al. 2017) squamates (Burbrink et al. 2020; Singhal et al. 2021), birds (Jarvis et al. 2014; Prum et al. 2015), and amphibians (Hime et al. 2021), but there has not yet been an investigation into how gene tree-species tree discordance across the genome affects the inference of relationships between the major turtle lineages. While phylogenetic conflict often belies more traditional means of measuring certainty via bootstrap support or posterior probabilities (Minh et al. 2020a), the study of turtle systematics has not yet truly taken advantage of the genomic era; this is despite ample genomic resources being available for a wide range of turtle species (Shaffer et al. 2013; Wang et al. 2013; Tollis et al. 2017; Quesada et al. 2019). Since evolutionary distinctiveness is linked to conservation status in turtles (Colston et al. 2020), and many turtle species have adaptations of physiological and developmental importance such as longevity, resistance to hypoxia, cold tolerance, and the anatomical changes responsible for the turtle shell, a study using genome-scale data to measure phylogenomic heterogeneity is needed in order to obtain the most accurate models of turtle evolution. Our goals in this study were threefold: (1) to revisit discoveries about higher turtle systematics using substantial proportions of the genome; (2) to measure heterogeneity at genes and sites in important events in turtle evolution; and (3) determine what drives heterogeneity in turtle phylogenomics in terms of mechanisms and locus type.

To accomplish these goals, we generated two distinct yet overlapping genomic datasets for turtles. The first comprises 5,310 high-confidence single-copy orthologs bioinformatically extracted from the genome assemblies of 19 turtles plus three outgroups, aligned and phylogenetically analyzed using multilocus coalescent-consistent and concatenated partition-based methods. The second dataset (26 species, including all turtle species from the first method) consisted of 1,655,675 biallelic parsimony-informative sites that we extracted using mapped Illumina sequence reads. We assembled the reads into contigs that were further mapped to a reference turtle genome (the western painted turtle, *Chrysemys picta bellii*). Based on genome annotations, we partitioned sites by locus type including coding regions (CDS), introns, 5’-UTR, 3’-UTR, intergenic, pseudogenes, lncRNA, and smRNA, and reconstructed separate phylogenies using each locus type. With the first dataset we assessed patterns and sources of heterogeneity in turtle phylogenomics with a large number of DNA sequence alignments, while with the second dataset we compared topologies reconstructed with parsimony-informative sites extracted from different annotated regions across the turtle genome. Taken together, this scope of analyses is unprecedented in turtle phylogenomics. We confirm that heterogeneity in phylogenomic datasets is to be expected, even in well-resolved clades such as turtles, and suggest that a combination of processes is driving the incongruence between previous studies of turtle relationships.

## Methods and Materials

### Phylogenomic analysis of high-confidence single-copy orthologs

We downloaded the complete genome assemblies of 19 turtle species plus three outgroup taxa from NCBI. The species, assembly accession numbers, assembly lengths, and assembly contiguities measured by scaffold N50 are shown in Table 1. Using BUSCO v3 (Waterhouse et al. 2018), we extracted the nucleotide sequences of 5,310 OrthoDB v9 tetrapod orthologs (Zdobnov et al. 2017) from each genome assembly. BUSCO orthologs are high-confidence genes that persist in eukaryotic genomes as single-copy, which precludes downstream problems in phylogenetics stemming from gene duplication such as hidden paralogy (Waterhouse et al. 2018). We aligned each set of orthologs with MAFFT v7.475 (Katoh and Standley 2013), and removed erroneous columns in the alignments with the heuristic method (*automated 1*) in TrimAL v1.4.1 (Capella-Gutierrez et al. 2009). Summary statistics including number of taxa, alignment length, missing data percentage, number and proportion of variable sites, number and proportion of parsimony-informative sites, and GC content for each alignment were estimated with AMAS v3.04 (Borowiec 2016). Trimmed alignments were filtered at a cut-off length ≥1,500 bp and a minimum of 50% taxa representation. We estimated the average pairwise Kimura 2-parameter corrected distance for each alignment with MEGAX (Kumar et al. 2018).

**Table 1.**
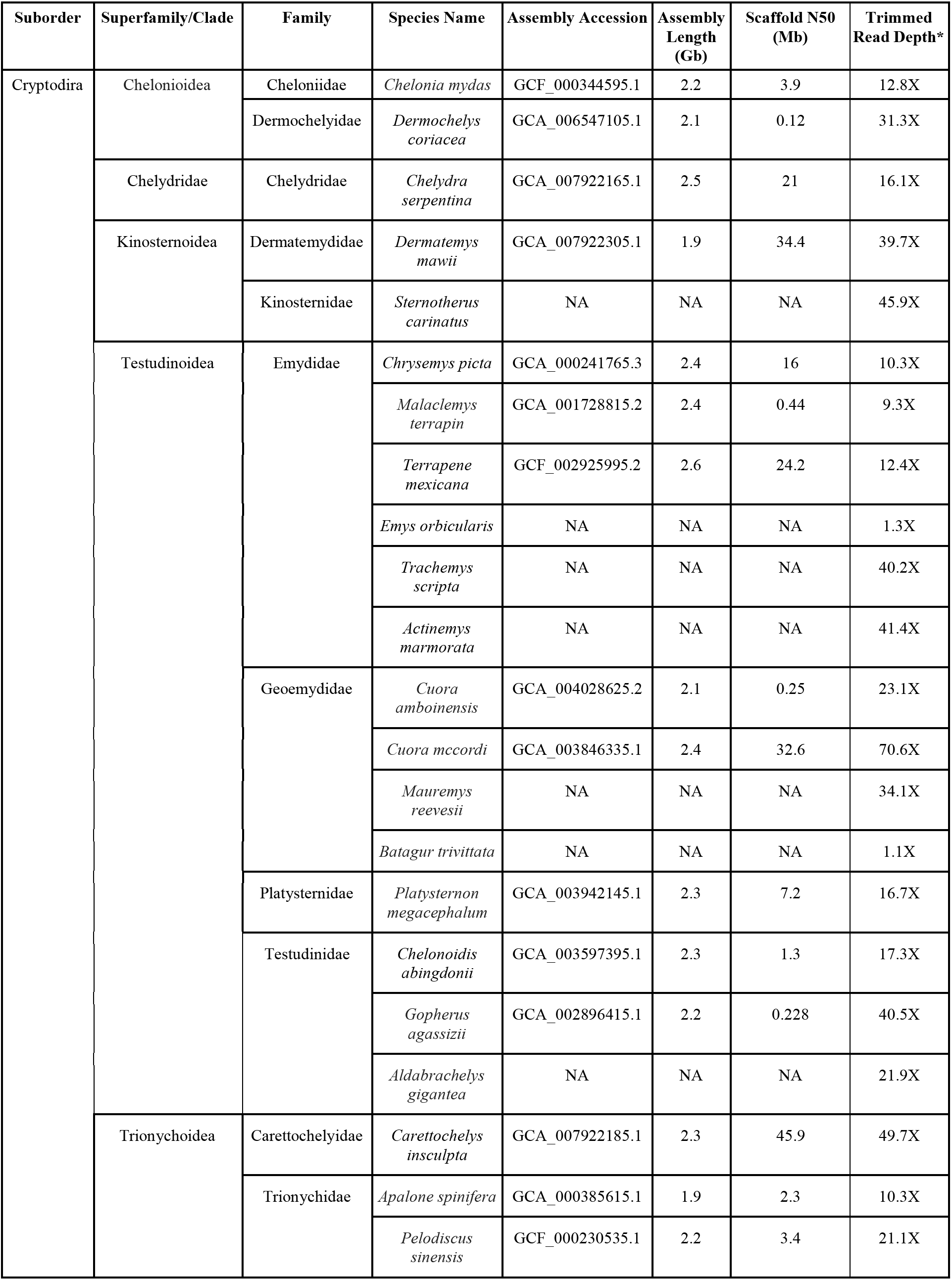
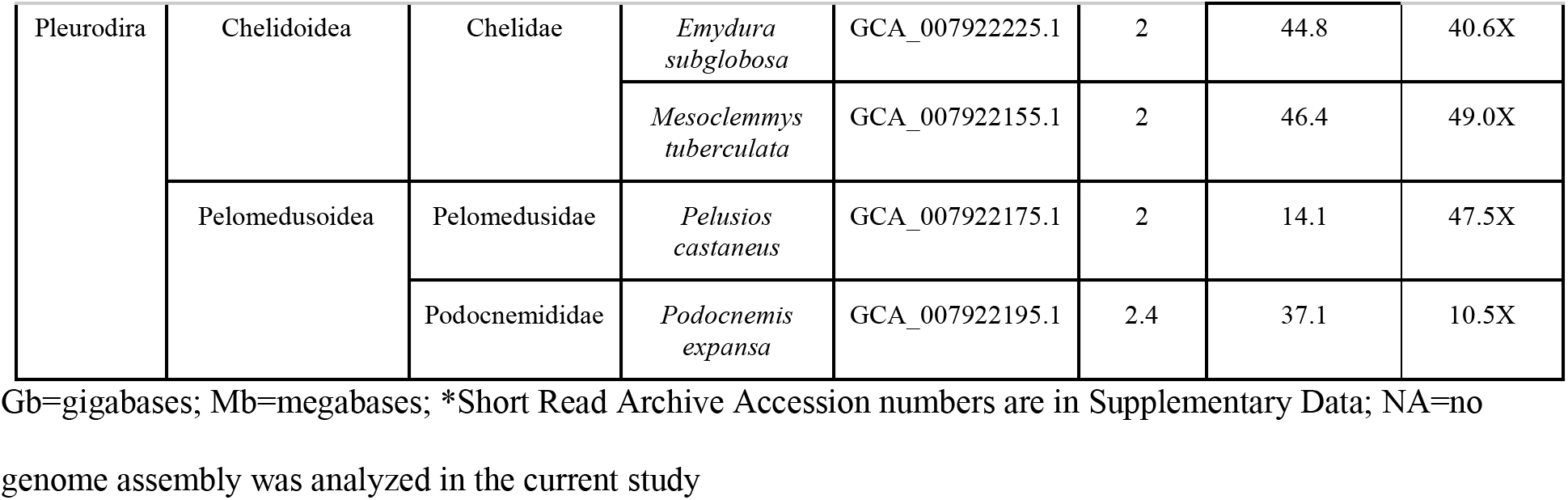
Genomic data for turtles analyzed in this study.

Gene trees were reconstructed for each locus using RAxML v1.8 (Stamatakis 2014). For each ortholog, we generated 10 maximum likelihood trees with the GTRGAMMA substitution model, and performed 100 rapid bootstrap replicates on the best tree. We then collected the best tree for each ortholog to use as evidence to construct a species tree with ASTRAL-III (Zhang et al. 2018), assessing branch support with local posterior probabilities (Sayyari and Mirarab 2016). We used Snakemake (Mölder et al. 2021) to automate a reproducible bioinformatics pipeline for BUSCO extraction, filtering, alignment, gene tree and species tree estimation, available at https://github.com/mibyars/busco_phylo_pipeline. We also concatenated the single copy orthologs into a supermatrix with AMAS and performed a partitioned maximum likelihood phylogenetic reconstruction with IQ-TREE v2.1.2 (Minh et al. 2020b). We used ModelFinder (Kalyaanamoorthy et al. 2017) set to TESTMERGE to combine partitions where necessary. We assessed branch support on the concatenated partitioned maximum likelihood tree using 1,000 ultrafast bootstrap replicates and the Shimodaira-Hasegawa-like (SH-like) approximate likelihood ratio test (aLRT, Shimodaira 2002).

### Phylogenomic analysis of biallelic markers

In addition to single-copy orthologs from turtle and outgroup genome assemblies, we constructed sets of biallelic single nucleotide polymorphisms (SNPs) using paired-end sequence data from multiple turtle species downloaded from the NCBI Short Read Archive (SRA, Table 1, Supplementary Data). We assessed each SRA dataset with FastQC v0.115 (S. Andrews - http://www.bioinformatics.babraham.ac.uk/projects/fastqc/) and trimmed the raw reads with BBDuk v.37.41 (B. Bushnell - sourceforge.net/projects/bbmap/, last accessed December 2020). We used SISRS to generate de novo orthologs from the processed reads (Schwartz et al. 2015; Literman and Schwartz 2021). After subsampling all sequences to a target depth of ∼10X per taxon (except for two low-coverage species, Table 1, Supplementary Data), we assembled a composite genome using Ray v2.2.3-devel (Boisvert et al. 2010) with default parameters and a k-value of 31. We then used SISRS to map the trimmed reads from each species to the composite genome, retaining only uniquely mapped reads to avoid false positives from duplicated or repetitive regions. Bases in the composite genome were replaced according to mapped data from each species if sites were covered by at least three reads and if there was no within-taxon variation; sites not meeting these criteria were masked as “N”.

For downstream ortholog annotation, we mapped each scaffold from the composite sequences against the reference genome for the western painted turtle (*Chrysemys picta bellii,* GCA_000241765.3) with Bowtie2 v2.3.4 (Langmead and Salzberg 2012), removing contigs that either failed to map or mapped to multiple positions, and removing sites resulting from overlapping coverage of independent scaffolds. We then annotated the mapped composite genome sites parsed from the painted turtle genome as belonging to the following genomic regions (locus types) with BEDTools v2.2.6 (Quinlan and Hall 2010): (1) coding sequences (CDS); (2) 3’-UTR; (3) 5’-UTR; (4) introns; (5) other genic regions not annotated as CDS, UTR, or intronic; (6) long-noncoding RNAs (lncRNAs); (7) pseudogenes; (8) smRNAs (including miRNAs, ncRNAs, rRNAs, scRNAs, smRNAs, snoRNAs, snRNAs, tRNAs, and vaultRNAs).; and (9) intergenic regions. We defined “intergenic” as all regions not included within the other locus types.

We used SISRS to identify parsimony-informative sites along the mapped composite contigs and to remove invariant sites and singletons, as well as any sites containing “Ns” (i.e. polymorphic within species) and overlapping indels or gaps. For the nine annotated genomic regions as well as a set of the complete mapped data (=10 locus types), we concatenated biallelic parsimony-informative sites and inferred locus type phylogenies via SVDquartets (Chifman and Kubatko 2014) in PAUP* v4.0a (build 168) (Swofford 2003) using 1,000 bootstrap replicates and estimated majority consensus trees.

### Heterogeneity analyses

To measure similarity across genealogies, we calculated Robinson-Foulds (RF) distances (Robinson and Foulds 1981) with PhyKIT (Steenwyk et al. 2021). First, we measured the RF distance between each inferred single-copy ortholog gene tree and the inferred species tree. In order to take into account missing species across some of the gene trees, we normalized the RF distances by taking the raw RF distance and dividing it by 2(*n*-3) where *n* is the number of tips in the phylogeny. We also calculated multiple pairwise RF distances between the trees inferred from SISRS data mapped to each genome annotation (CDS, introns, intergenic, etc.). Finally, we calculated the RF distances of each SISRS phylogeny from the topology inferred with all mapped sites across the turtle genome.

We computed gene and site concordance factors for each branch in the turtle phylogeny using the trimmed and length-filtered single-copy ortholog alignments, their gene trees, and the ASTRAL tree with IQ-TREE (Minh et al. 2020a). A branch’s gene concordance factor represents the percentage of decisive gene trees that also contain that branch, and a branch’s site concordance factor is the percentage of decisive alignment sites supporting that branch. We also computed gene concordance factor and site concordance factor for each branch in the concatenated single-copy ortholog tree by computing individual locus trees based on the partitions in IQ-TREE. Locus trees were inferred with default settings, and the species tree was inferred using an edge-linked proportional partition model with an SH-like approximate likelihood ratio test and ultrafast bootstrap analysis of 10,000 replicates each (-alrt 10000 -B 10000). The greedy algorithm of PartitionFinder (Lanfear et al. 2012) was applied to find the best-fit partitioning scheme which was then used in the subsequent tree reconstruction step (-m TESTMERGE). We also calculated site concordance factors for every branch in the SVDquartets trees inferred from biallelic parsimony-informative sites within locus types.

Discordance between gene trees and species trees can be caused by introgression, incomplete lineage sorting, erroneously inferred gene trees stemming from model violations and/or other artifacts, or combinations of all of these factors, and we aimed to determine how these issues affect phylogenomic inference in turtles. One expectation of incomplete lineage sorting is a roughly equal proportion of topologies supporting alternative relationships, while introgression would cause some minor gene trees to be more frequent than others, reflecting the direction of gene flow (Huson et al. 2005; Green et al. 2010). We collected the two most common minor topologies and each node (gDF1 and gDF2 resulting from the concordance factor analysis in IQ-TREE) and used a chi-squared test to determine if they were similar in terms of their proportions of all gene trees, where a rejection of the null hypothesis was interpreted as the presence of introgression in addition to incomplete lineage sorting and potential model violations. Nodes where we failed to reject the null hypothesis were interpreted as divergence patterns in turtle evolution driven primarily by either incomplete lineage sorting or model violations.

### Analysis of potential model violations via substitutional saturation, compositional heterogeneity, and codon usage bias

Phylogenetic signal can be obscured by the accumulation of multiple substitutions at the same site over time, leading to model violations and increased gene tree-species tree discordance, particularly at deep evolutionary timescales (Jeffroy et al. 2006; Philippe et al. 2011). Therefore, to further distinguish between the effects of incomplete lineage sorting and model violations at nodes in the turtle species tree, we estimated levels of substitutional saturation in the single-copy ortholog dataset. First, we calculated the slope of a linear regression between the computed raw pairwise distances and corrected pairwise distances under a TN93 substitution model for 685 single-copy ortholog alignments with complete taxon sampling using the ape 5.5 R package (Paradis and Schliep 2019). For genes with high levels of saturation, the slope of this regression will be closer to 0 while less saturated genes will have a slope approaching 1 (Philippe et al. 1994). We then plotted a histogram of all slopes and characterized each alignment as having slopes above (“unsaturated”) and below (“saturated”) the mean. To determine the effect of substitutional saturation on phylogenomic inference in turtles, we compared ASTRAL species trees inferred from the RAxML trees inferred from unsaturated and saturated genes.

To analyze substitutional saturation at codon positions and assess their effects on phylogenomic inference in turtles, we realigned the 685 single-copy orthologs with complete taxon sampling using MACSE (Ranwez et al. 2011), and concatenated alignments that were partitioned based on (1) first and second codon positions and (2) third codon positions with AMAS. We then computed the corrected and uncorrected pairwise distances and estimated the regression slopes for the two codon-based partitions in R as above. We also further separated these codon partitions according to whether they belonged to the unsaturated or saturated gene sets, inferred maximum likelihood phylogenies on the four resulting datasets in IQ-TREE with model testing and 1,000 ultrafast bootstrap replicates, and calculated site concordance factors for each inferred tree from codon partitions. Finally, to account for substitutional saturation we computed a maximum likelihood tree from the concatenated and partitioned MACSE amino acid alignments of the 685 single-copy orthologs using model testing in IQ-TREE. Maximum likelihood tree topologies, branch support, and concordance factors from codon-partitioned saturated and unsaturated alignments as well as the amino acids were compared to the ASTRAL and concatenated supermatrix trees based on single-copy orthologs ≥1,500bp.

A chi-squared test for lineage-specific compositional heterogeneity is computed before each IQ-TREE run, where the frequency of each nucleotide in a given taxon is compared to the overall frequency of each nucleotide in the entire dataset. We compared the results of this test across IQ-TREE analyses of the saturated and unsaturated concatenated partitions based on codon position. To analyze codon usage bias in our single copy ortholog dataset, we estimated how often a codon is observed relative to its predicted occurrence in the absence of codon usage bias, by computing the Relative Synonymous Codon Usage (RSCU) of the concatenated 685 complete-taxon MACSE alignments using CodonW v1.4.4 (Peden 1999) followed by factor analysis in FactoMineR (Lê et al. 2008) in R.

### Divergence time estimation

We estimated divergence times with single-copy ortholog alignments using Bayesian modeling implemented in BEAST v.2.6.3 (Bouckaert et al. 2019). We subsampled 685 alignments with complete taxon sampling, from which we further randomly sampled three replicate sets of 10 genes. Based on model testing with ModelFinder, we applied the HKY+Gamma+Invariant site model to all partitions, and utilized a relaxed lognormal molecular clock and Calibrated Yule tree prior. To calibrate the time trees, we used 11 calibration priors from the literature (Joyce et al. 2013) which overlapped with our taxon sampling (Supplementary Data). To account for uncertainty in fossil dating, we used hard minimum constraints and set soft maximum constraints by placing the maximum ages in the 97.5% quantile of a lognormal prior distribution. Markov chain Monte Carlo (MCMC) analyses were sampled every 50,000 generations. For each replicate, we assessed convergence of parameter estimates across the MCMC by monitoring effective sample sizes with Tracer v.1.7.2 (Rambaut et al. 2018), and we estimated a maximum clade credibility tree with TreeAnnotator v2.6.3.

We also estimated divergence times in turtle evolution using penalized likelihood with r8s v1.8.1 (Sanderson 1997, 2002, 2003), using the same fossil calibrations as above for minimum and maximum node ages; the main difference in this analysis was we fixed the time to most recent common ancestor (TMRCA) for amniotes at 312 million years ago (Ma) (Donoghue and Benton 2007) and the TMRCA for Testudines at 220 Ma (Thomson et al. 2021). First, we concatenated the 685 complete partitions and inferred a maximum likelihood tree with model testing for each partition as above in IQ-TREE. We then used the inferred phylogeny with branches in terms of substitutions per site and the 1,836,182 concatenated bases for penalized likelihood estimation, with cross validation to optimize the smoothing parameter which quantifies the deviation from the molecular clock.

We used linear regression to determine the relationship between the estimated divergence time for each node and the concordance factors computed with the inferred trees from each locus type in the SISRS analysis, the gene trees and the species tree based on single-copy orthologs, and the locus trees and partitioned maximum likelihood tree based on single-copy orthologs. We also tested for a correlation between the concordance factors and the estimated rates of molecular evolution at each branch. We repeated these regressions using estimated divergence times from Thomson et al. (2020).

## Results

### A very large number of informative sites for turtle phylogenomics

We analyzed publicly available genome assemblies to extract single copy orthologs from 19 turtles representing 13 out of 14 extant families and 7 of 8 superfamilies, plus 3 outgroup taxa (Table 1). On average, 4,609 (83%, range 1,911-5,099) conserved tetrapod orthologs were complete in the genome assemblies, 112 (2.1%, range 7-496) were duplicated, 170 (3.2%, range 55-485) were fragmented, and 419 (7.9%, range 143-2,907) were missing (Fig. 1). Fifty-five percent of conserved tetrapod orthologs were missing from the genome assembly of the trionychid *Apalone spinifera*; therefore, we omitted this species from downstream analyses of single-copy orthologs due to concerns about assembly quality (Waterhouse et al. 2018). Each single-copy ortholog was present in an average of 19 out of 21 species, averaged 1,902 bp in length, and contained an average of 53% variable sites and 26% parsimony-informative sites. Alignment length was correlated with the number of parsimony-informative sites (R^2^=0.69, P=2.2e-16, Fig. 2a). GC content was consistent across the single-copy orthologs and averaged 48% (standard deviation 6.2%, Fig. 2b), as was average pairwise Kimura 2-parameter distance (mean 0.154, median 01.39, standard deviation 0.072, Fig. 2c). The subset of 2,513 alignments ≥1,500 bp in length had very similar characteristics to the complete set of loci; the average large locus length was 1,903 bp and the relationship between alignment length and the number of parsimony-informative sites held (R^2^=0.73, P=2.2e-16).

**Figure 1.**
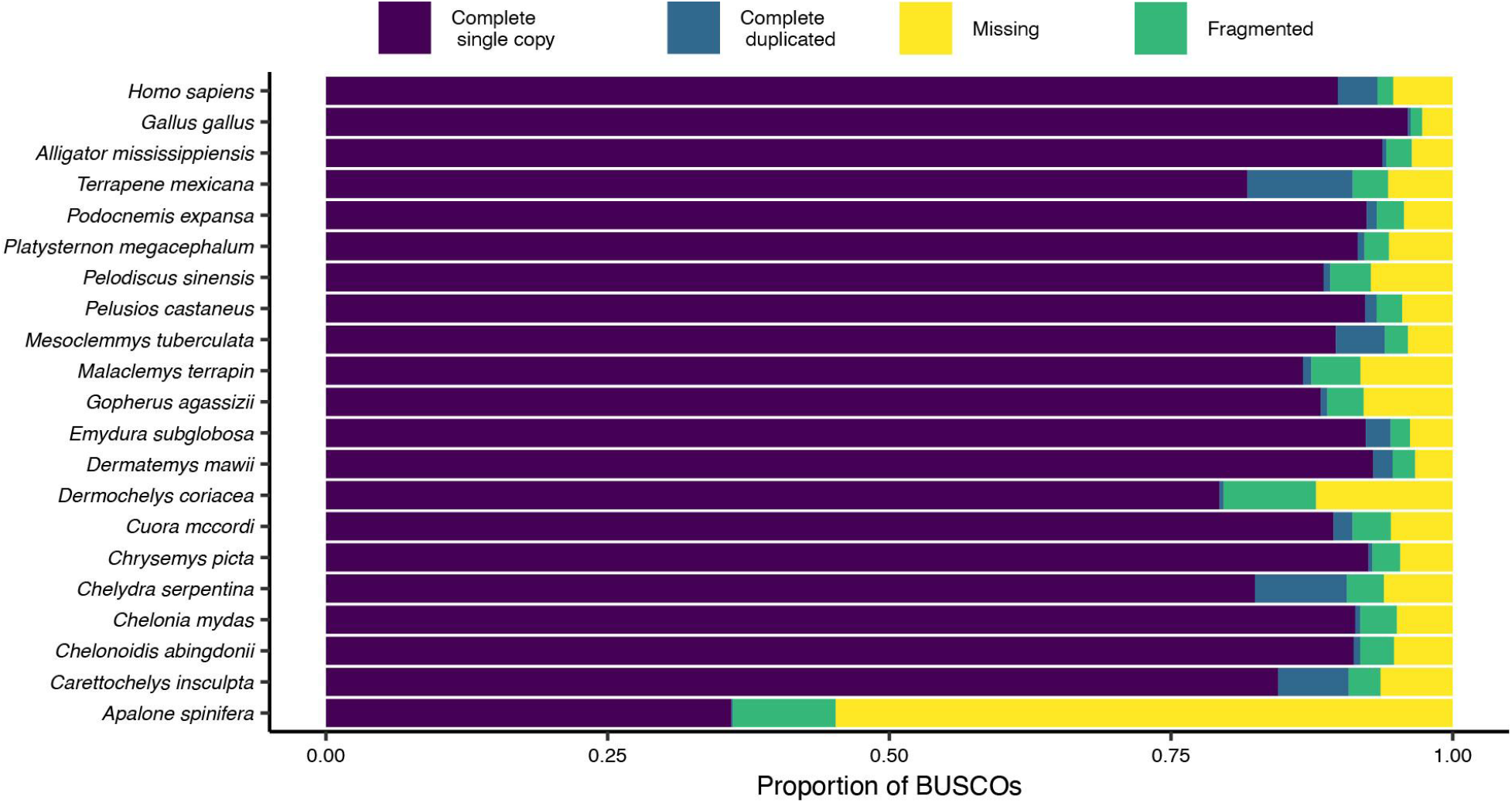
Summary of single-copy orthologs. Percent of tetrapod orthologs from orthoDBv9 present in each genome assembly as complete single copy, complete duplicated, fragmented, or missing.

**Figure 2.**
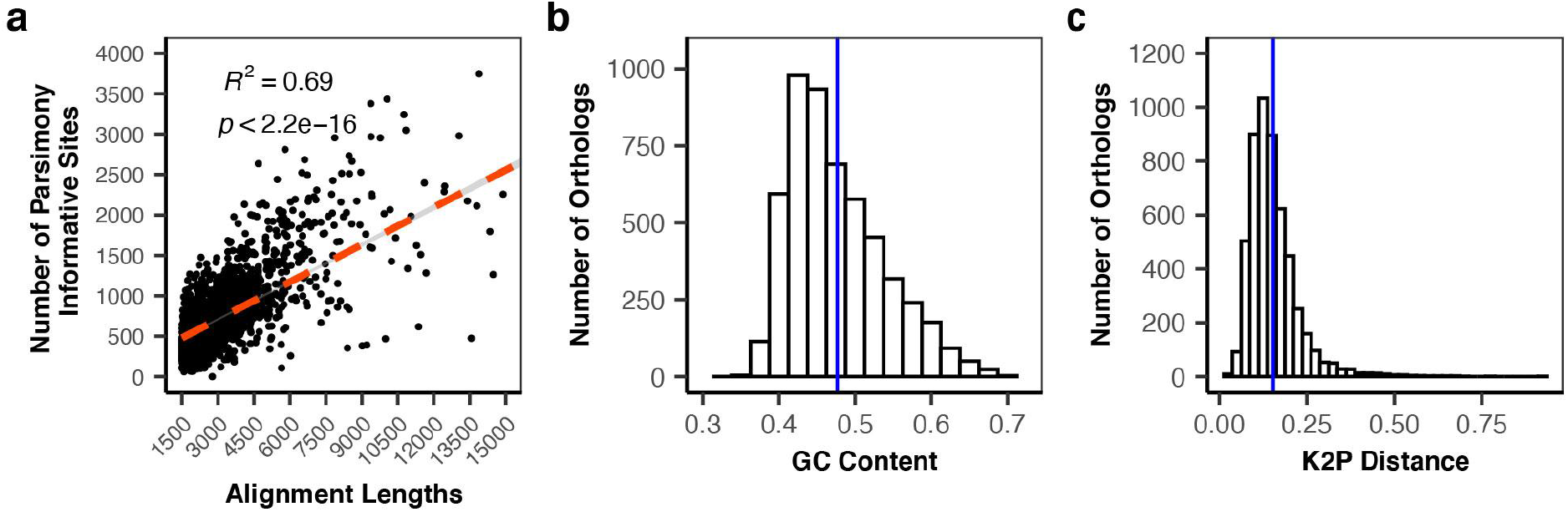
Description of single copy ortholog alignments. (a) Scatterplot showing the relationship between alignment length and the number of parsimony-informative sites for trimmed alignments ≥1,500 bp. (b) Histogram showing the distribution of GC content in all single copy orthologs after trimming. Solid vertical line represents the mean value (0.476). (c) Histogram showing the distribution of average pairwise Kimura 2-parameter (K2P) distances for single copy orthologs after trimming. Solid vertical line represents the mean value (0.154).

We downloaded 38 SRA datasets from 26 turtle species representing 7 of 8 superfamilies and 14 families (Table 1; Supplementary Data), including the 19 species with complete genome assemblies from the first dataset. Based on FastQC results, we selected 32 of the SRA datasets for further analysis ranging from ∼1–70X post-trimming coverage. The composite genome consisted of 5,555,666 contigs with an N50 of 2.4 Mb. We successfully mapped 302,355,876 bases to 2,110,354 contigs in the composite genome which covered 80% of the painted turtle genome (Table 2), and called 1,655,675 parsimony-informative biallelic SNPs when allowing missing data for up to two species at a given site. The number of parsimony-informative biallelic SNPs was much larger in mapped datasets containing only cryptodires (2,787,072 bases) and testudinoideans (4,696,637 bases) (Supplementary Data), as expected based on Literman et al. (2021). We collected the highest number of biallelic SNPs from intergenic regions (746,724 bases), followed by introns (533,336 bases) and CDS (136,823 bases). The relatively low number of parsimony-informative biallelic SNPs in pseudogenic (427) and smRNA regions (137) is likely due to a small number of annotations for these features in the western painted turtle genome.

**Table 2.** Summary of short read data from 26 turtle species mapped to the western painted turtle (*Chrysemys picta*) genome annotations.

| Locus Type | Annotated Bases in the Genome | Bases Mapped by SISRS | Number of biallelic parsimony-informative sites* | Percent of Annotated Bases Mapped |
| --- | --- | --- | --- | --- |
| All Mapped | 2,365,766,571 | 302,355,876 | 1,655,675 | 12.78% |
| Intergenic | 1,382,145,112 | 161,445,195 | 746,724 | 11.68% |
| Intronic | 812,010,672 | 115,297,781 | 533,336 | 14.20% |
| lncRNA | 64,450,450 | 9,451,244 | 52,207 | 14.66% |
| Genic Other | 65,764,053 | 9,052,113 | 36,604 | 13.76% |
| CDS | 33,486,790 | 5,718,074 | 136,823 | 17.08% |
| 3'-UTR | 8,034,514 | 1,510,783 | 15,150 | 18.80% |
| 5'-UTR | 5,096,255 | 708,214 | 6,519 | 13.90% |
| Pseudogenic | 670,023 | 59,058 | 427 | 8.81% |
| smRNA | 133,894 | 20,076 | 137 | 14.99% |

\*When allowing up to two missing species per site, see Supplementary Data for numbers of sites with varying levels of missing data.

### Bioinformatic analysis of turtle genomes yielded well-resolved species relationships

The inferred relationships among turtle lineages based on single-copy orthologs were consistent with previous studies using phylogenomic markers (Crawford et al. 2015; Shaffer et al. 2017), with full statistical support for all resulting branches based on posterior probabilities in the ASTRAL tree, and bootstrap replicates and SH-like approximate likelihood in the supermatrix tree (Fig. 3, Supplementary Fig. 1, Supplementary Fig. 2). Hereafter we refer to these results collectively as the turtle species tree. The turtle species tree supports the reciprocal monophyly of Pleurodira and Cryptodira (Fig. 4). Within cryptodires, there is a split between the Trionychoidea (soft-shelled turtles) and the Durocryptodira (Crawford et al. 2015), and a further subdivision of Durocryptodira into the Testudinoidea (pond turtles, big-headed turtles, geoemyds, and tortoises) and Americhelydia (sea turtles, snapping turtles, mud turtles, and *Dermatemys*; Joyce et al. 2013). Within the Testudinoidea, the turtle species tree supports Testuguria (geoemyds and tortoises; Joyce et al. 2004), and a *Platysternon* + Emydidae clade (“Platyemidae”; Barley et al. 2010; Crawford et al. 2015).

**Figure 3.**
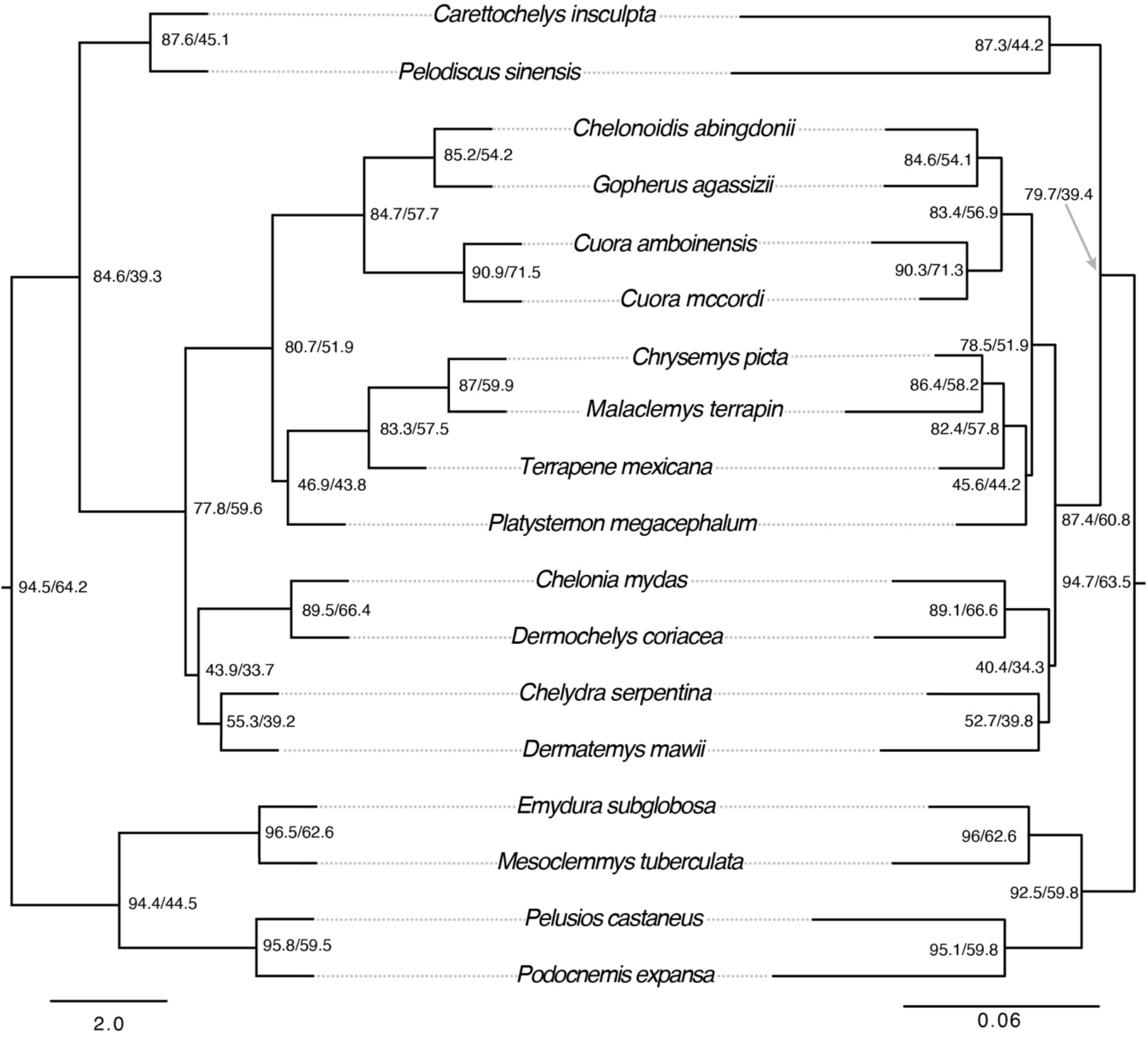
Species trees for turtles based on high-confidence single-copy orthologs. We used 2,513 single-copy orthologs ≥1,500 bp in length to infer a multilocus coalescent-consistent tree based on gene genealogies (ASTRAL, left) and a maximum likelihood tree (ML, right) from a concatenated and partitioned supermatrix in IQ-TREE. All branches received 100% support from local posterior probabilities (ASTRAL), 1,000 bootstrap replicates (IQ-TREE), and approximate likelihood ratio test (IQ-TREE). Node labels indicate gene and site concordance factors before and after the slash, respectively. Branch lengths are in terms of coalescent units (ASTRAL) and substitutions per site (ML). Outgroups (*Gallus, Alligator, Homo*) are not shown.

**Figure 4.**
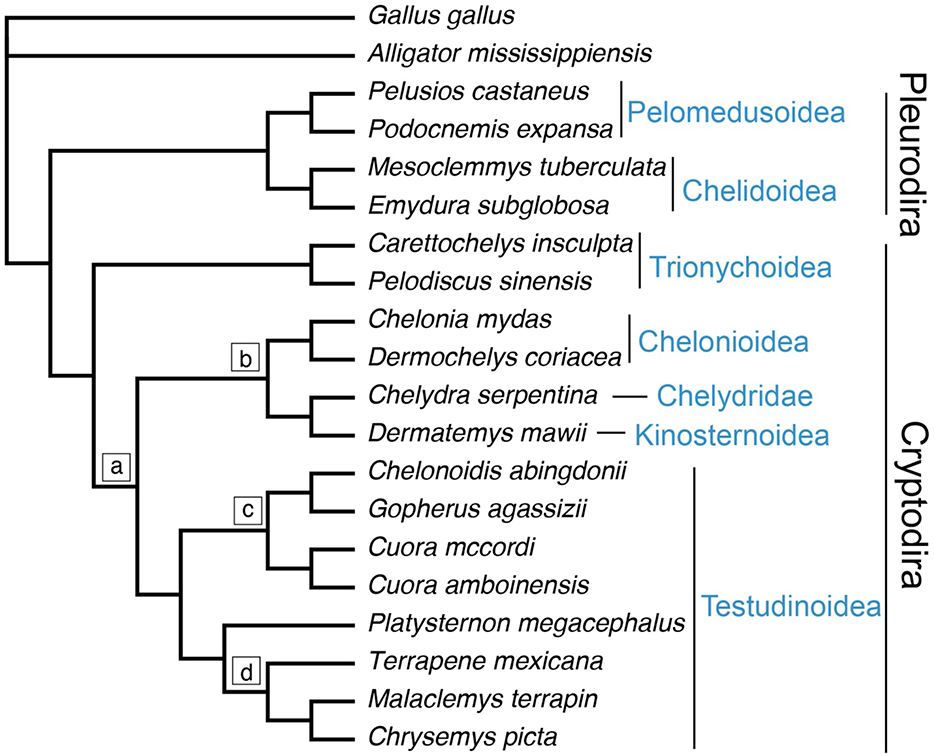
The phylogenetic relationships of higher turtle taxa. Cladogram depicting the turtle species tree in agreement with results from ASTRAL and concatenated partitioned maximum likelihood analyses as well as most SVDquartets analyses using the mapped data (although these analyses lacked outgroups). Superfamilies or clades are indicated by horizontal shaded text. Additional key clades are labelled with letters in boxes. (a) Durocryptodira, (b) Americhelydia, (c) Testuguria, (d) Emydidae. Additional outgroup *Homo sapiens* not shown.

SVDquartets trees from the genome-wide, all-mapped data as well as from CDS, introns, and intergenic regions were consistent with the species tree based on single-copy orthologs (Fig. 4, Supplementary Fig. 3-11; Supplementary Data). When there was disagreement between trees from mapped regions and the species tree, the difference was often in the placement of *Platysternon* (Fig. 5). For instance, when allowing zero missing gapless parsimony-informative sites from the all-turtle data, 5’ UTR placed *Platysternon* as the outgroup of Geoemydidae + Testudinidae (435 sites). When allowing up to two missing gapless parsimony-informative sites from the all-turtle data, pseudogenic regions (426 sites) also placed *Platysternon* as the outgroup of Geoemydidae + Testudinidae, while smRNA (137 sites) agreed with the species tree and all-mapped topologies. Meanwhile, at multiple levels of allowed missing data the Testudinoidea-only mapped sites weakly supported *Platysternon* as the sister taxon to Geoemydidae + Testudinidae + Emydidae (Supplementary Data).

**Figure 5.**
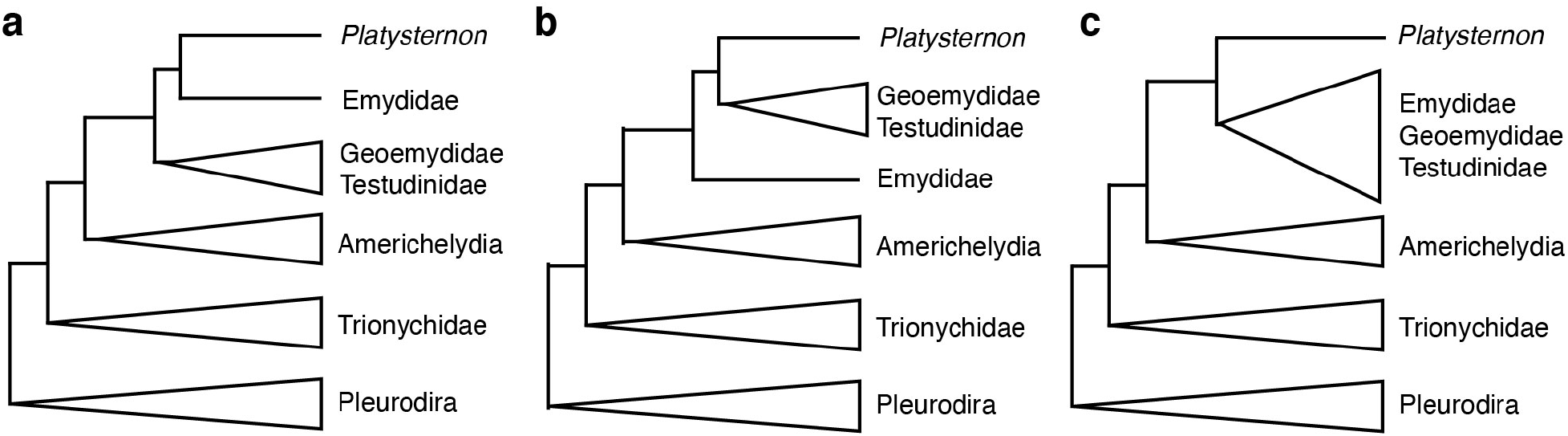
Alternative topologies in turtle phylogenetics based on markers from different regions of the genome. (a) *Platysternon* as the sister taxon to Emydidae to the exclusion of Testuguria (Geoemydidae + Testudinidae), consistent with the turtle species tree (current study), Crawford et al. (2015), and Shaffer et al. (2017). (b) *Platysternon* as the sister taxon to Testuguria with the exclusion of Emydidae, consistent with 17% of the single-copy ortholog genes trees, mapped 5’-UTR parsimony-informative sites data with zero missing data, mapped pseudogene parsimony-informative sites when allowing 2 missing taxa per site, and Krenz et al. (2005). (c) *Platysternon* as the sister taxon to Geoemydidae + Testudinidae + Emydidae, consistent with 20% of single-copy ortholog gene trees and Shaffer et al. (1997).

### Heterogeneity is prevalent at well resolved nodes in the turtle phylogeny

While results from our single-copy orthologs and biallelic sites analyses of turtles are consistent and largely in agreement with other studies based on 100% branch support (Crawford et al. 2015; Shaffer et al. 2017), some genomic regions, either from single-copy orthologs or from genome feature annotations, yielded genealogies conflicting with the turtle species tree. For instance, based on normalized RF distances most gene trees from single-copy orthologs were similar to the species tree, although there was a long tail to this distribution indicating gene trees with greater dissimilarity (Fig. 6a). SVDquartets trees using mapped data from CDS, introns, and intergenic regions were consistent with both the single-copy ortholog species tree and a SVDquartets tree derived from the all-mapped data. In contrast, mapped pseudogenic DNA, 5’ UTR, and smRNA deviated from the all-mapped data in terms of their RF distances (Supplementary Fig. 12).

**Figure 6.**
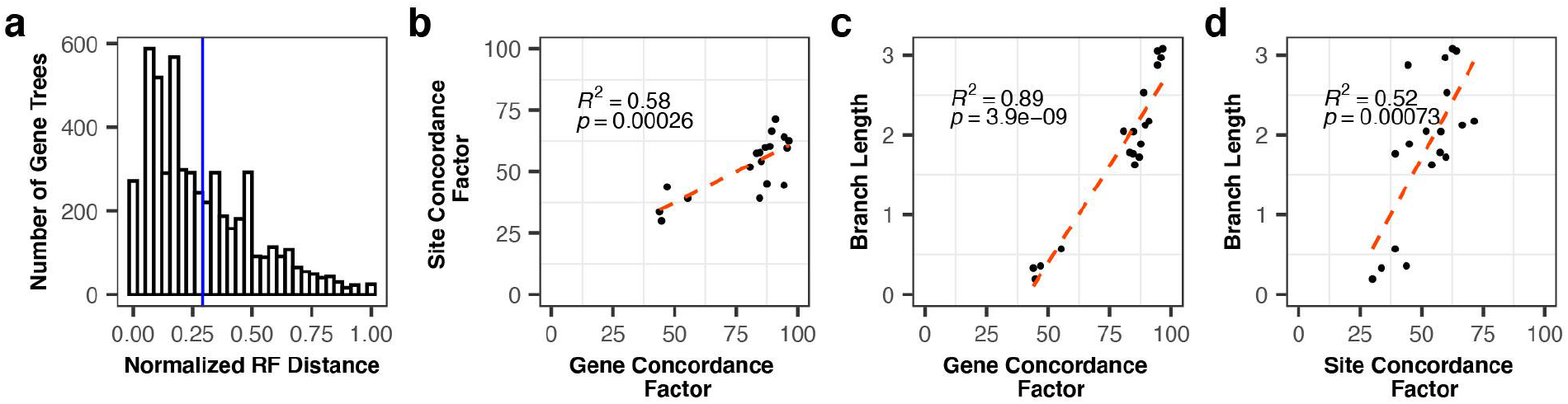
Phylogenomic heterogeneity in turtles. (a) Histogram of RF distances between gene trees and species tree. (b-c) Concordance factors calculated using single-copy ortholog alignments, their gene trees, and the ASTRAL species tree. Results based on locus trees and concatenated supermatrix maximum likelihood tree are in the supplement. (b) Scatterplot depicting the relationship between gene and site concordance factors at each node. (c) Scatterplot depicting the relationship between gene concordance factor and branch length. (d) Scatterplot depicting the relationship between site concordance factor and branch length.

We also found a high degree of heterogeneity based on gene and site concordance factors throughout the turtle phylogeny (Fig. 3, Fig. 6, Supplementary Fig. 1, Supplementary Fig. 2). For instance, while local posterior probabilities were uniformly 100% across branches of the ASTRAL tree, gene concordance factors ranged from 44 to 97 and site concordance factors ranged from 30 to 71 (Fig. 6b-d). Gene and site concordance factors based on the ASTRAL tree were correlated with each other (Figure 6b; R^2^=0.55, P=0.0002) and with branch lengths (Fig. 5c-d; R^2^=0.88, 3.88e-09 and R^2^=0.49, P=0.0007, respectively). We also estimated gene and site concordance factors based on the locus trees with respect to the maximum likelihood tree inferred from the concatenated supermatrix, and found a similar range of gene concordance factors (34 to 92) and site concordance factors (30 to 71) despite 100% bootstrap support, as well as a strong correlation between the two concordance measures (R^2^=0.60, P=0.0001). Gene and site concordance factors based on the concatenated partitioned maximum likelihood tree were also correlated with branch lengths (R^2^=0.90, 1.02e-09 and R^2^=0.58, P=0.0001, respectively).

The ancestral branch leading to turtles was assigned a relatively large gene concordance factor and site concordance factor across analyses based on the ASTRAL, supermatrix, and SVDquartets trees, and there was high concordance for reciprocally monophyletic Pleurodira and Cryptodira across genes and sites (Fig. 3, Fig. 4, Supplementary Figs. 3-10). The placement of Trionychoidea as sister taxon to all other cryptodires was assigned a relatively large gene concordance factor but small site concordance factor, suggesting that there are a large number of sites supporting alternative bifurcations. One area with particularly high discordance (i.e. relatively small gene concordance factor and site concordance factor across all datasets) was the placement of the big-headed turtle (*Platysternon*) with respect to Emydidae. Our chi-squared test rejected introgression at this node (Table 3), favoring incomplete lineage sorting or potential model violations as sources of discordance in the position of *Platysternon*. There was also a high degree of discordance associated with the americhelydian taxa (Chelonioidea + Chelydridae + Kinosternoidea), particularly with the ancestral branch leading to Americhelydia as well as with the sister taxon relationship between *Dermatemys* (Kinosternoidea) and *Chelydra* (Chelydridae). Based on single-copy orthologs, minor gene tree frequencies were significantly different at these nodes (Table 4), suggesting that introgression in addition to incomplete lineage sorting and/or model violations may be driving phylogenomic discordance in these groups.

**Table 3.**
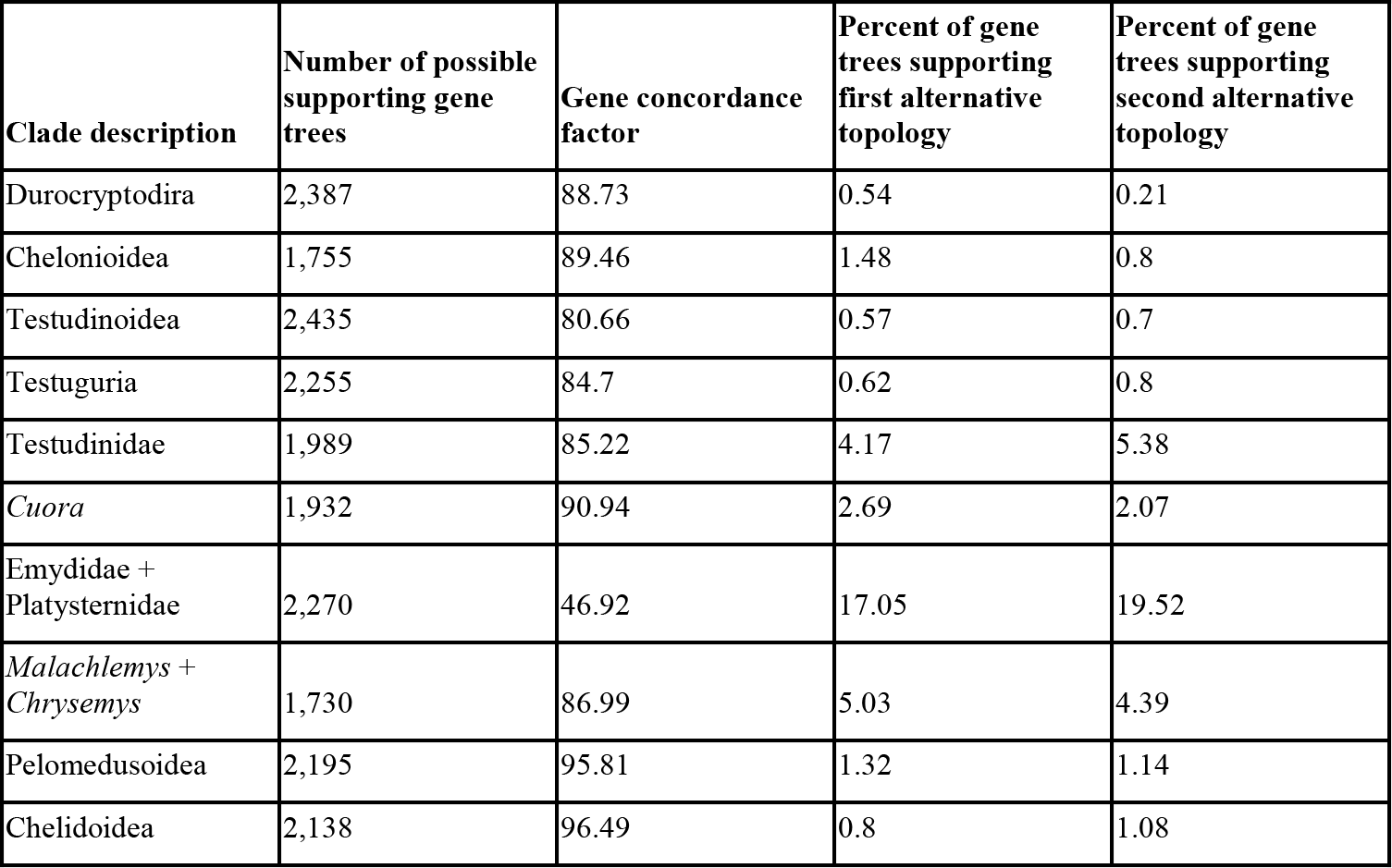
Gene concordance factors for clades represented by nodes in the ASTRAL species tree phylogeny where the minor gene tree frequencies were similar according to a chi-squared test.

**Table 4.**
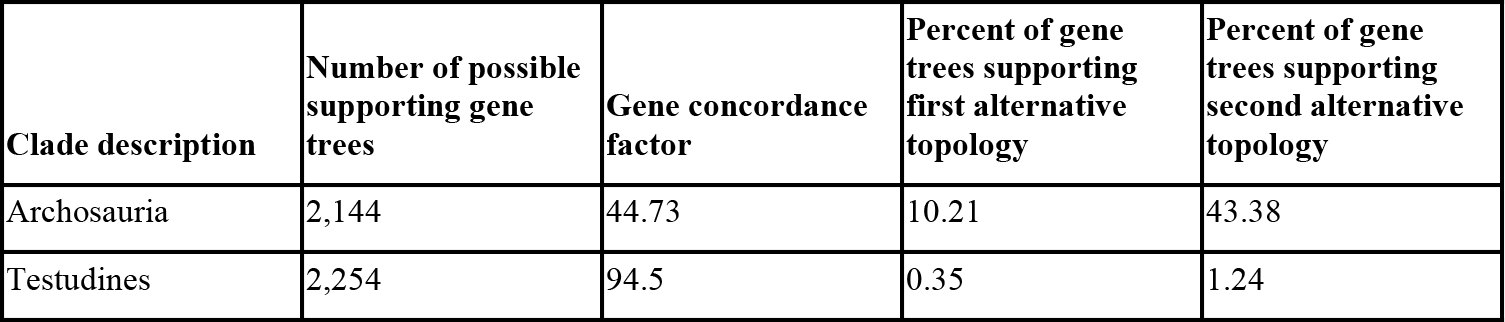
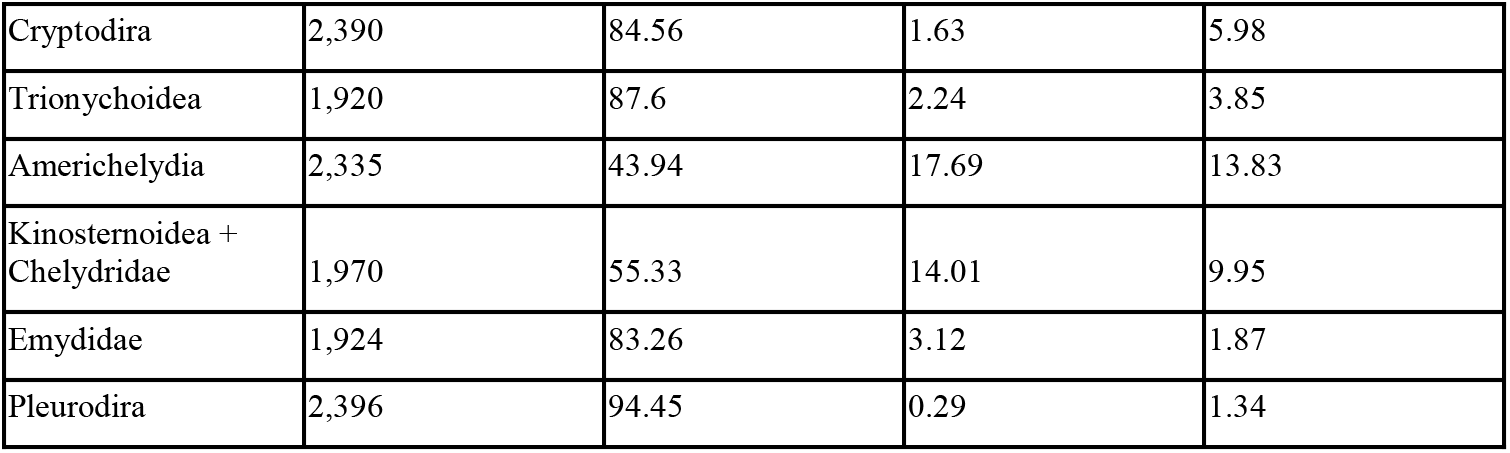
Gene concordance factors for clades represented by nodes in the ASTRAL species tree phylogeny where the minor gene tree frequencies were statistically different according to a chi-squared test.

| Clade description | Number of possible supporting gene trees | Gene concordance factor | Percent of gene trees supporting first alternative topology | Percent of gene trees supporting second alternative topology |
| --- | --- | --- | --- | --- |
| Archosauria | 2,144 | 44.73 | 10.21 | 43.38 |
| Testudines | 2,254 | 94.5 | 0.35 | 1.24 |
| Cryptodira | 2,390 | 84.56 | 1.63 | 5.98 |
| Trionychoidea | 1,920 | 87.6 | 2.24 | 3.85 |
| Americhelydia | 2,335 | 43.94 | 17.69 | 13.83 |
| Kinosternoidea + Chelydridae | 1,970 | 55.33 | 14.01 | 9.95 |
| Emydidae | 1,924 | 83.26 | 3.12 | 1.87 |
| Pleurodira | 2,396 | 94.45 | 0.29 | 1.34 |

### Patterns of substitutional saturation, compositional heterogeneity, and codon usage bias in turtle phylogenomics

We measured the slope of the line in a regression between uncorrected and corrected genetic distances for 685 single-copy ortholog alignments with complete taxon sampling. The slopes ranged from *y*=0.47*x* (most saturated) to *y*=0.91*x* (least saturated), but were skewed towards low levels of saturation, with a mean of *y*=0.76*x*, a first quartile of *y*=0.72*x*, and a third quartile of *y*=0.81*x* (Supplementary Fig. 13). When analyzed by codon position, we found that first and second codon positions were unsaturated (*y*=0.86*x*) while the slope of the line between corrected and uncorrected distances at third codon positions were consistent with saturation (*y*=0.57*x*) (Supplementary Fig. 14). Saturation did not have an effect on the topology of the species tree, as ASTRAL trees derived from the most and least saturated sets of genes were identical (Supplementary Data). In addition, all codon partitions regardless of their saturation levels produced maximum likelihood trees with intra-Testudines relationships that were identical in topology to the species tree, and the maximum likelihood tree estimated based on amino acid partitions was also identical in topology to the species tree (Supplementary Data). There was some disagreement in the placement of turtles in the amniote phylogeny that appeared to be driven by substitutional saturation; for instance, the maximum likelihood trees based on first and second codon positions from saturated genes and all sets of third codon positions supported turtles as the sister taxon to crocodilians (Supplementary Data; see Discussion).

We found that compositional heterogeneity was most prominent in the saturated third position and first and second position datasets, as well as in the saturated first and second position dataset (Supplementary Data). The unsaturated alignment of first and second codon positions exhibited the least amount of compositional heterogeneity, with five species differing significantly from the rest of the alignment; however, these species were distributed randomly across the turtle species tree. Correspondence analyses of the RSCU revealed that some codons, particularly CCG (Proline), ACG (Threonine), and GCG (Alanine) were favored in different species (Supplementary Fig. 15). However, there was little evidence for a phylogenetic signal in this pattern within turtles since codon usage bias was dispersed randomly among species, and the only outlier along the second dimension was an outgroup taxon (*Gallus*).

### Heterogeneity at different timescales in turtle evolution

Results from the replicate BEAST analyses consistently underestimated turtle divergence times with respect to results from the penalized likelihood analysis as well as those of Thomson et al. (2020) (Supplementary Data), although are closer to estimates from Chiari et al. (2012) and Pereira et al. (2017). Across locus types from the SISRS analyses, we found that site concordance factors were not strongly correlated with estimated node ages (Fig. 7, Supplementary Fig. 16). For most locus types the relationship between site concordance factor and divergence time at a given node trended slightly negatively, suggesting a pattern of higher concordance at younger nodes, although these comparisons were non-significant (Fig. 7b, P>0.1). The one exception to this trend was for pseudogenes, for which the relationship between site concordance factors and estimated divergence times trended positive albeit without significance or with only moderate significance (P=0.072 using Thomson et al. 2020’s divergence time estimates). The relationship between site concordance factors and estimated DNA substitution rates trended slightly positive for all locus types from the SISRS analysis, but was non-significant in all comparisons (Supplementary Fig. 17).

**Figure 7.**
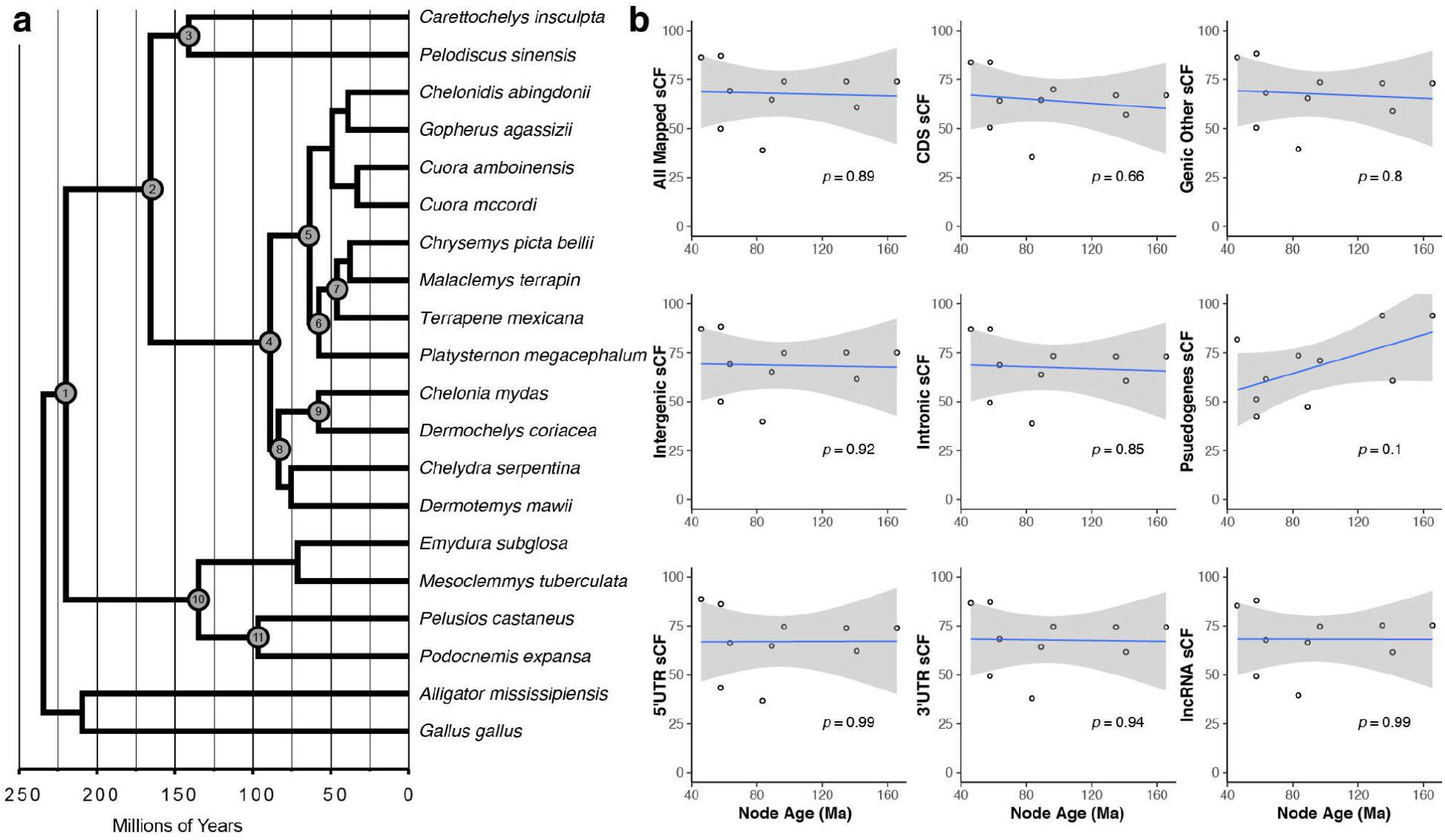
Site concordance and estimated node ages in turtle evolution. (a) Chronogram showing estimated divergence times in millions of years based on penalized likelihood and fossil calibrations of labelled nodes. 1. Testudines, 2. Cryptodira, 3. Trionychoidea, 4. Durocryptodira, 5. Geoemydidae, 6. Platyemidae, 7. Emydidae, 8. Americhelydia, 9. Chelonioidea, 10. Pleurodira, 11. Pelomedusoidea. (b) Site concordance factors (sCF) for each node in phylogenies inferred from nine locus types and the estimated node age in millions of years ago (Ma) based on penalized likelihood.

We also found a nonsignificant negative trend in the relationship between site concordance factor and divergence time at a given node based on both the ASTRAL and supermatrix trees based on single-copy orthologs (Fig. 8a-c; Supplementary Fig. 18. However, we found a positive (albeit non-significant) trend in the relationship between the estimated divergence time at a given node and its gene concordance factor when using both trees derived from single-copy orthologs (Fig. 8e-g; Supplementary Fig. 18). There was no clear relationship between the site concordance factor of a given node and the estimated substitution rate at the preceding branch (Fig. 8d), and a moderately significant positive trend (P≤0.05) for the relationship between the gene concordance factor of a given node and the estimated substitution rate at the preceding branch (Fig. 8h; Supplementary Fig. 18).

**Figure 8.**
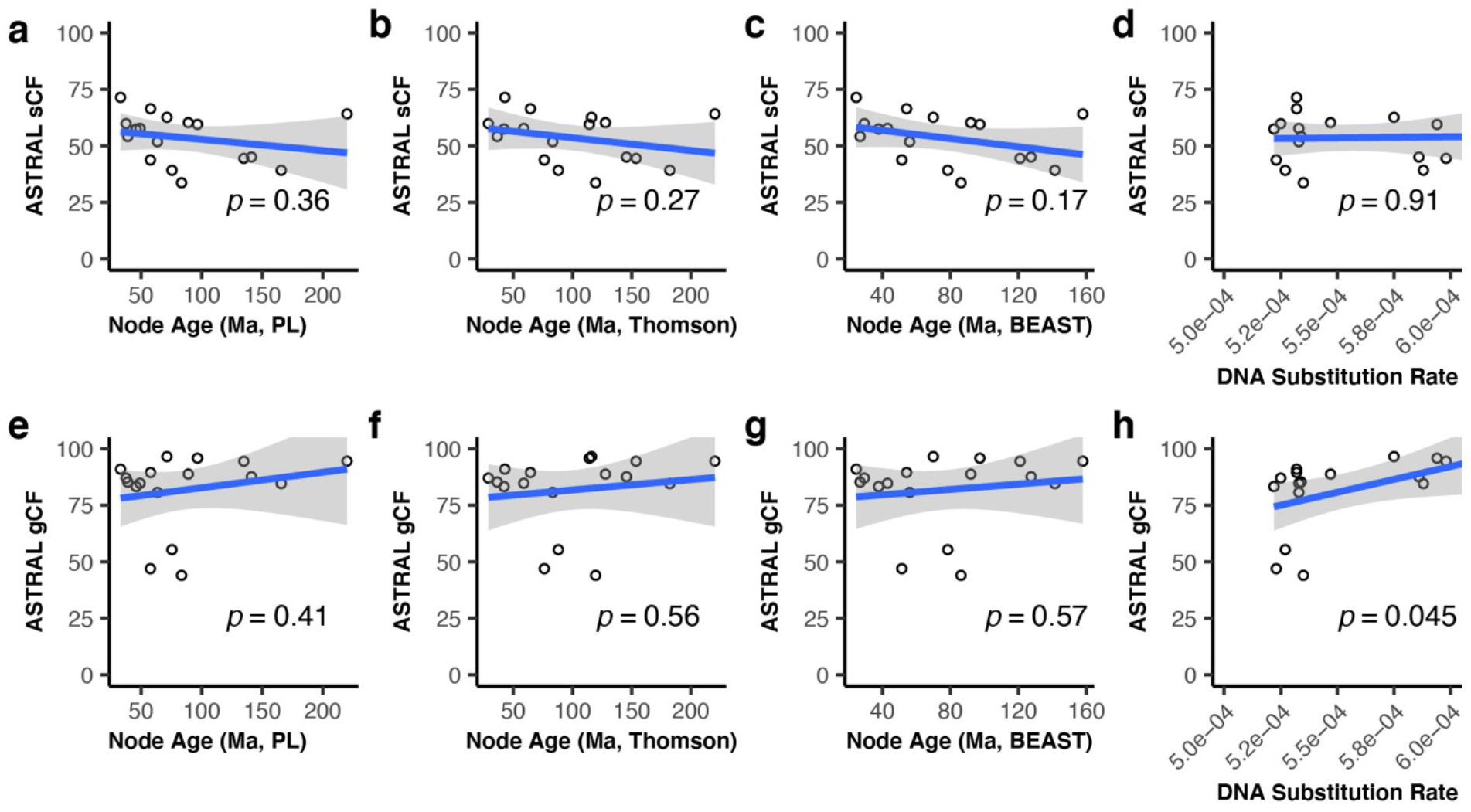
Concordance at genes and sites, estimated node ages, and DNA substitution rates in turtle evolution. Results shown are from comparisons of single-copy ortholog alignments and their gene trees to the species tree inferred using ASTRAL. There was a negative trend between site concordance factors (sCF) and estimated node age (in millions of years or Ma) based on penalized likelihood (PL) from the current study (a), estimated node ages in Ma from Thomson et al. (2020) (b), estimated node ages in Ma from the BEAST analysis in the current study (c), and no discernable trend between sCF and estimated DNA substitution rates in terms of substitutions per site per Ma (d). There was a positive trend between gene concordance factors (gCF) and estimated node ages (e, f, g) and a moderately significant positive relationship between gene concordance factors and estimated DNA substitution rates (h). Results from comparisons of the locus trees and the maximum likelihood tree inferred from the concatenated supermatrix are shown in Supplementary Materials.

## Discussion

Heterogeneity has been recognized as an important feature of phylogenomic datasets, and the exploration of the patterns and sources of gene tree and site discordance has become an essential step when reconstructing the branching order of life with molecular data (Kumar et al. 2012; Minh et al. 2020a). To our knowledge, while the role of gene tree-species tree discordance in estimating the position of turtles within the amniote phylogeny has been the focus of a few studies (Chiari et al. 2012), there has not been an assessment of phylogenomic heterogeneity and its effects on the assessment of intra-turtle relationships until now. Here, we have constructed a robust phylogenomic dataset for turtles based on whole genomes in order to sample the major turtle lineages. We identified splits in the turtle phylogeny and regions of the turtle genome associated with gene tree and site heterogeneity, and investigated how potential sources of this discordance such as introgression, incomplete lineage sorting, and technical issues leading to model violations may have contributed to conflicting results in phylogenetic estimation of turtles. The results of our study demonstrate that heterogeneity can be rampant, even in phylogenomic datasets which have yielded fully resolved species trees.

Protein-coding genes such as the single-copy orthologs used in the current study are common markers used for phylogenetics, due to their straightforward amplification via PCR or RNA-Seq, as well as relatively straightforward methods for their orthology assignment and alignment. However, the use of coding sequences in phylogenetics is complicated by difficulties with accurate evolutionary modeling due to multiple hits at ancient divergences, a diversity of substitution rates across the genome, and selective constraint potentially limiting the number of variant sites. These problems often scale with the size of phylogenomic datasets and can lead to biased results (Philippe et al. 2011). Therefore, in addition to the single-copy orthologs we also generated a complementary dataset for turtle phylogenomics consisting of millions of parsimony-informative biallelic sites using SISRS. Employment of these sites has multiple advantages: (1) they are easily extracted from multiple sequence alignments; (2) they do not require complicated evolutionary modeling (i.e. they either support or do not support a splitting event); and (3) the sheer number of potential sites allows strict quality filtering without loss of signal (Literman et al. 2021). SISRS resulted in a considerable number of sites collected from both coding, noncoding, and regulatory parts of the genome, including intergenic regions which yielded almost 750,000 parsimony-informative sites for turtles. These locus types sample a wide range of substitution rates, whereas many other phylogenomic methods focus on a much smaller range of locus types (such as UCEs, AHE, or coding genes). Our results show that many different regions of the turtle genome yield consistent support for the turtle species tree, leaving a solid phylogenetic hypothesis for higher turtle systematics. However, inferences from some regions such as pseudogenes and smRNA were based on a relatively small number of parsimony-informative sites, highlighting the need for more comprehensive annotations of noncoding portions of turtle genomes.

The relationships between turtles and other reptiles has been a longstanding debate in evolutionary biology, due to conflicting results between studies that use fossils and molecules (Benton et al. 2005; Chiari et al. 2012; Brown and Thomson 2017). Paleontological studies have placed turtles either at the root of the amniote phylogeny, or as sister taxa with various other reptilian groups, some of which are long extinct (Gaffney 1980). In the genomic era, analyses of nucleotides and protein sequences have been mostly in agreement, placing turtles as the sister taxon to archosaurs (birds and crocodilians) (Shaffer et al. 2013; Wang et al. 2013). Based on 2,513 long single-copy orthologs, our ASTRAL tree results supported a turtles + archosaurs clade, while the maximum likelihood tree based on the concatenated and partitioned supermatrix supported a turtles + crocodilians clade that excludes birds. Several lines of evidence suggest that the latter result is due to poorly fit evolutionary models. First, the maximum likelihood phylogeny inferred using partitioned amino acids, which accounts for saturation since proteins are less degenerate than nucleotides, yielded the turtles + archosaurs relationship. Second, we found that first and second codon positions from non-saturated genes supported turtles + archosaurs, while first and second codon positions from saturated genes as well as all third codon positions agreed with turtles + crocodilians (Supplementary Data). Brown and Thomson (2017) found that duplicated genes in the dataset published by Chiari et al. (2012) dataset were more likely to reject a turtles + archosaurs clade. Meanwhile, our reliance on single-copy orthologs from OrthoDB greatly reduces the role of paralogy in the conflict at this node. We suggest that the correct placement of turtles within the amniote phylogeny using genomic data has been hampered by substitutional saturation at such ancient divergences, and that when accounting for these artifacts the data support a turtles + archosaurs clade (Archelosauria; see Crawford et al. 2015).

When gene tree-species tree discordance is driven largely by technical errors such as saturation, it should be most apparent at older nodes in phylogeny, which are more affected by homoplasy, poor alignments, or model misspecifications. However, with the exception of the turtle-archosaur split, across both single-copy ortholog and SISRS datasets we found no significant correlation between concordance factors and estimated node ages. A slightly negative relationship between site concordance factors calculated from the single-copy ortholog dataset and node age suggests that these deeper splits in turtle evolution might be affected by multiple substitutions; however, this pattern is obscured a slightly positive relationship between gene concordance factors and node age using the same data. In addition, inferred turtle relationships were identical when analyzing different sets of genes separated by levels of substitutional saturation and partitioned by codon position. Thus, while it may be possible that multiple substitutions or other model violations are driving discordance at ancient nodes in the turtle phylogeny, their effects are either extremely weak or nonexistent within and among most turtle lineages, and the observed patterns may have more to do with errors in rate estimation using molecular data and fossil calibrations.

Our results suggest that population-level processes such as introgression and/or incomplete lineage sorting have played a key role at multiple stages of turtle diversification, leading to confusion in the literature about the placement of at least some taxa. In particular, the phylogenetic position of *Platysternon* has been highly contentious (Joyce et al. 2004, 2021), with early molecular studies in disagreement (Shaffer et al. 1997; Krenz et al. 2005; Parham et al. 2006; Barley et al. 2010). Our turtle species tree results mirror Crawford et al. (2015) and Shaffer et al. (2017), fully supporting *Platysternon* as the sister taxon to Emydidae. However, we found that only 47% of single-copy ortholog gene trees are in agreement with a *Platysternon* + Emydidae clade, and 37% of the remaining gene trees support only two alternative relationships: of these, 17% of gene trees support *Platysternon* as the sister taxon to Testuguria (Geoemydidae + Testudinidae) following Krenz et al. (2005), and 20% support *Platysternon* as the sister taxon to Testudinoidea (Emydidae + Geoemydidae + Testudinidae) following Shaffer et al. (1997). The proportion of gene trees supporting these alternative topologies was similar according to our chi-squared test (Table 3), and when we analyzed gene trees according to their levels of substitutional saturation, we saw no effect on the placement of *Platysternon*. As the most frequent alternate gene trees mirror published alternative hypotheses concerning the relationship of *Platysternon* to other testudinoideans, we suggest that incomplete lineage sorting is a previously unrecognized driver of conflicting phylogenetic results in the higher classification of these groups.

Another subject of conflict across previous phylogenetic studies of turtles is the relationship between Chelonioidea, Kinosternoidea, and the Chelydridae, all of which are now considered to be in the monophyletic Americhelydia (Barley et al. 2010; Joyce et al. 2013; Crawford et al. 2015). We found consistent support for Americhelydia across datasets and methods, as well as a sister taxon relationship between Chelydridae and Kinosternoidea, yet with considerable gene tree-species tree discordance with single-copy orthologs (Fig. 3) and site discordance with SISRS data (Supplementary Fig. 3-11). For instance, only 44% of decisive gene trees from single-copy orthologs contained an Americhelydia clade, and only 55% of decisive gene trees supported Chelydridae + Kinosternoidea. As with *Platysternon*, comparison of trees when accounting for saturation or codon position did not affect the placement of americhelydians. The frequencies of minor gene trees disagreeing with these splits were statistically different (Table 4), suggesting both introgression and incomplete lineage sorting may have played a role in the diversification of these taxa. To our knowledge, the role of hybridization during the speciation events at the deeper splits of the turtle phylogeny has not been investigated, although contemporary hybridization between sea turtle genera with up to ∼65 Ma of divergence is well documented (Karl et al. 1995; Arantes et al. 2020). As these occurrences are associated with low reproductive success (Arantes et al. 2020), it will be important to determine if episodic introgression is a feature of the long-term evolution of turtles.

Some recent molecular analyses of turtle relationships are limited in terms of the size of the molecular dataset, yet have more complete taxonomic sampling than our study, including up to 85% of extant species (Pereira et al. 2017; Colston et al. 2020; Thomson et al. 2021). A denser sampling of lineages maximizes the molecular signal for testing hypotheses about past diversification events as well as the relationship between evolutionary distinctiveness and extinction risk. In contrast, our study is limited in terms of taxonomic sampling due to our reliance on complete genomic sequences and assemblies for turtles, which are lagging compared to other amniote groups. As our results demonstrate widespread discordance in turtle phylogenomic datasets, additional lineages should be targeted for genomic sequencing in order to improve our ability to accurately estimate the evolutionary distinctiveness of certain species. These studies will be crucial in setting future conservation priorities for turtles.

## Acknowledgements

This research was funded by the National Science Foundation (#1812291 to RL) and by startup funds provided to MT by the School of Informatics, Computing, and Cyber Systems at Northern Arizona University. We would like to acknowledge the Monsoon computing cluster at Northern Arizona University (https://nau.edu/high-performance-computing/) for providing the computational resources necessary to carry out this study.

## Data Availability

The bioinformatic workflows used to extract and analyze single-copy orthologs from genome assemblies are available at https://github.com/mibyars/busco_phylo_pipeline. Data generated for this study including alignments and trees are available at the Harvard Dataverse (https://doi.org/10.7910/DVN/WYQRPC).

