## Supplementary Materials for "A Genomic Perspective on the Evolutionary Diversification of Turtles"

Simone M. Gable[1*], Michael I. Byars[1*], Robert Literman[2], Marc Tollis[1]

*[1]School of Informatics, Computing, and Cyber Systems, Northern Arizona University, PO Box 5693, Flagstaff, AZ 86011*

[2] *Department of Biological Sciences, University of Rhode Island, 120 Flagg Road, Kingstown, RI, 02881*

*These authors contributed equally.

**Supplementary Figure 1.** ASTRAL species tree based on single-copy orthologs. Branch lengths are in terms of coalescent units. Node labels are local posterior probabilities.

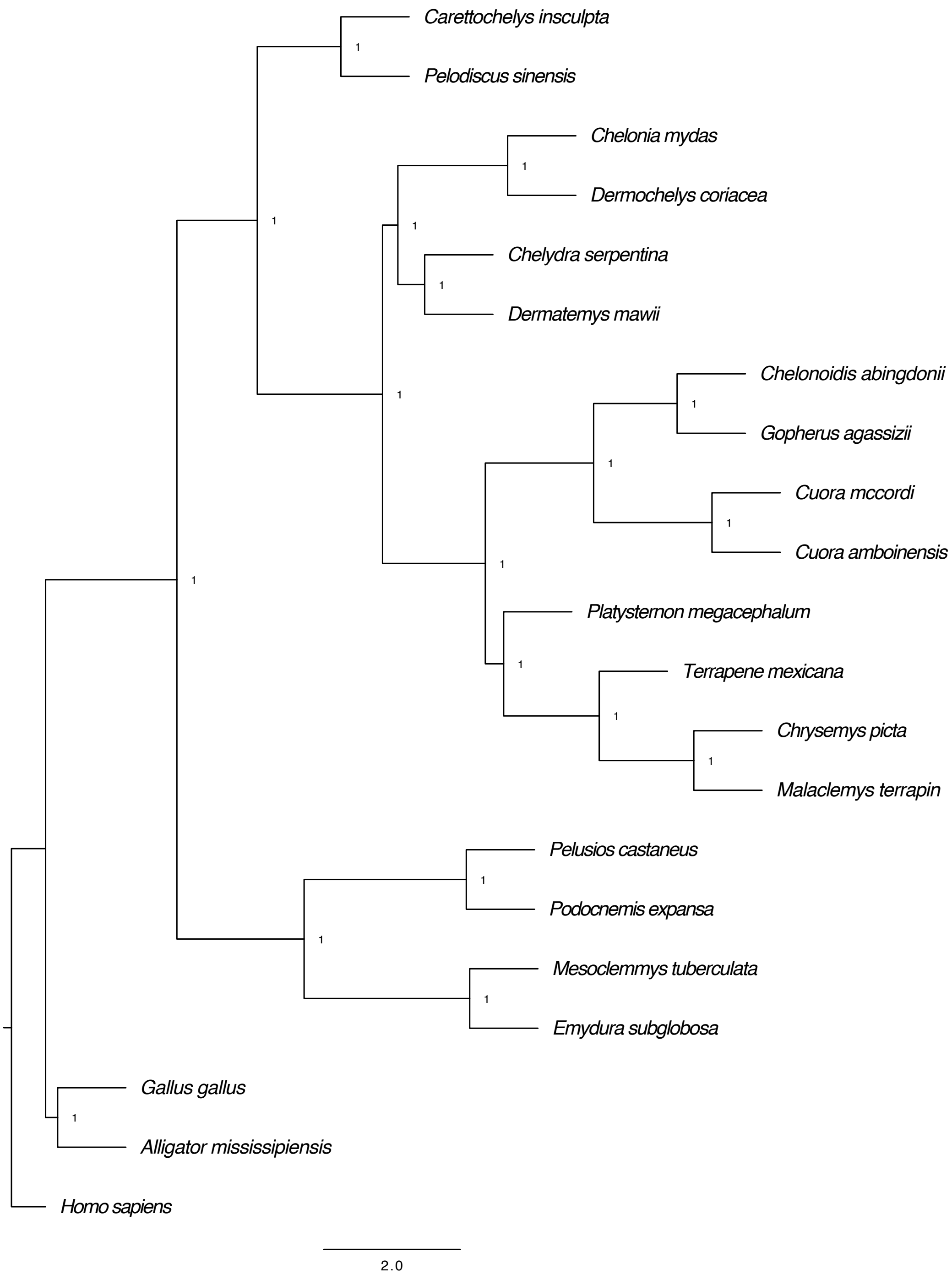

**Supplementary Figure 2.** Maximum likelihood tree based on concatenated and partitioned supermatrix of single-copy orthologs. Node labels are bootstrap support/approximate likelihood ratio test support/gene concordance factor/site concordance factor. Branch lengths are in terms of substitutions per site.

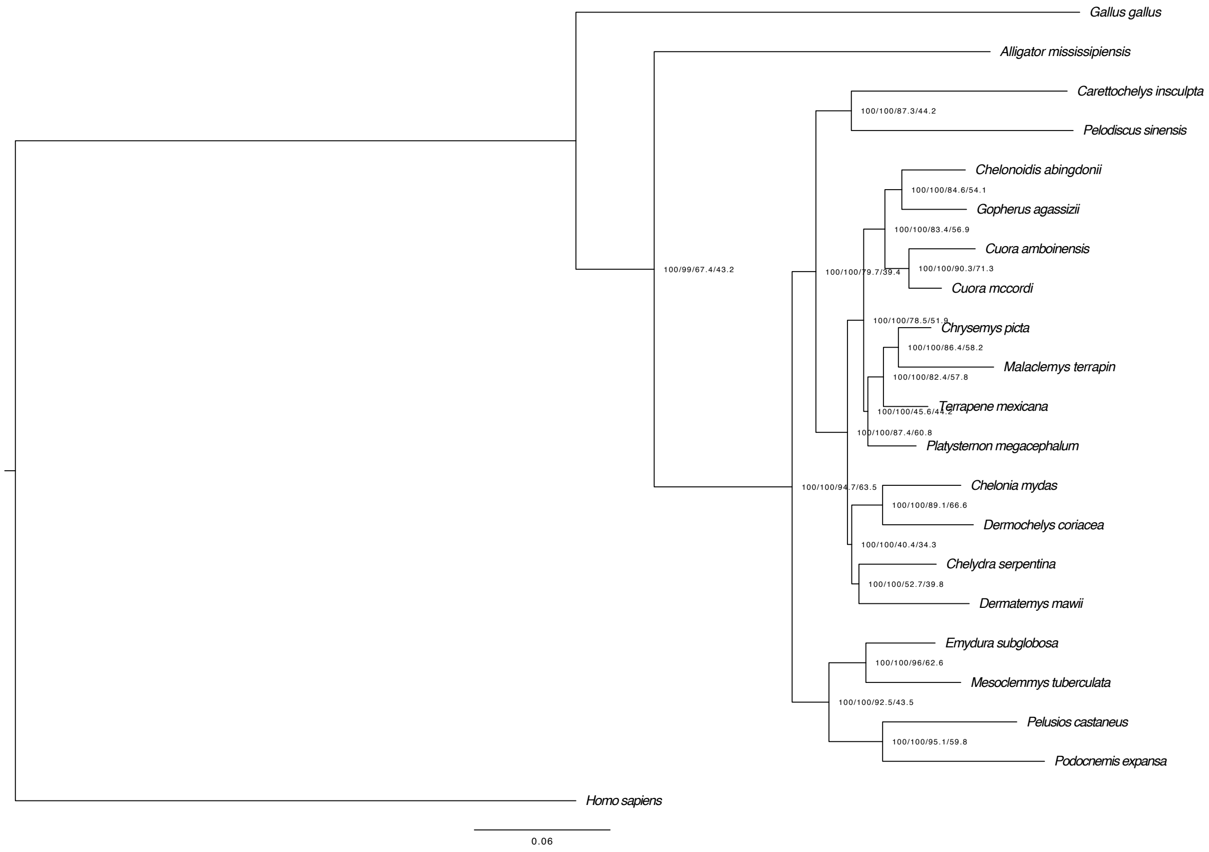

**Supplementary Figure 3.** SVDQuartets trees based on parsimony informative biallelic sites (All_Mapped). Top: support values represent the proportion of 100 bootstrap replicates. Bottom: Site concordance factors for each node are given.

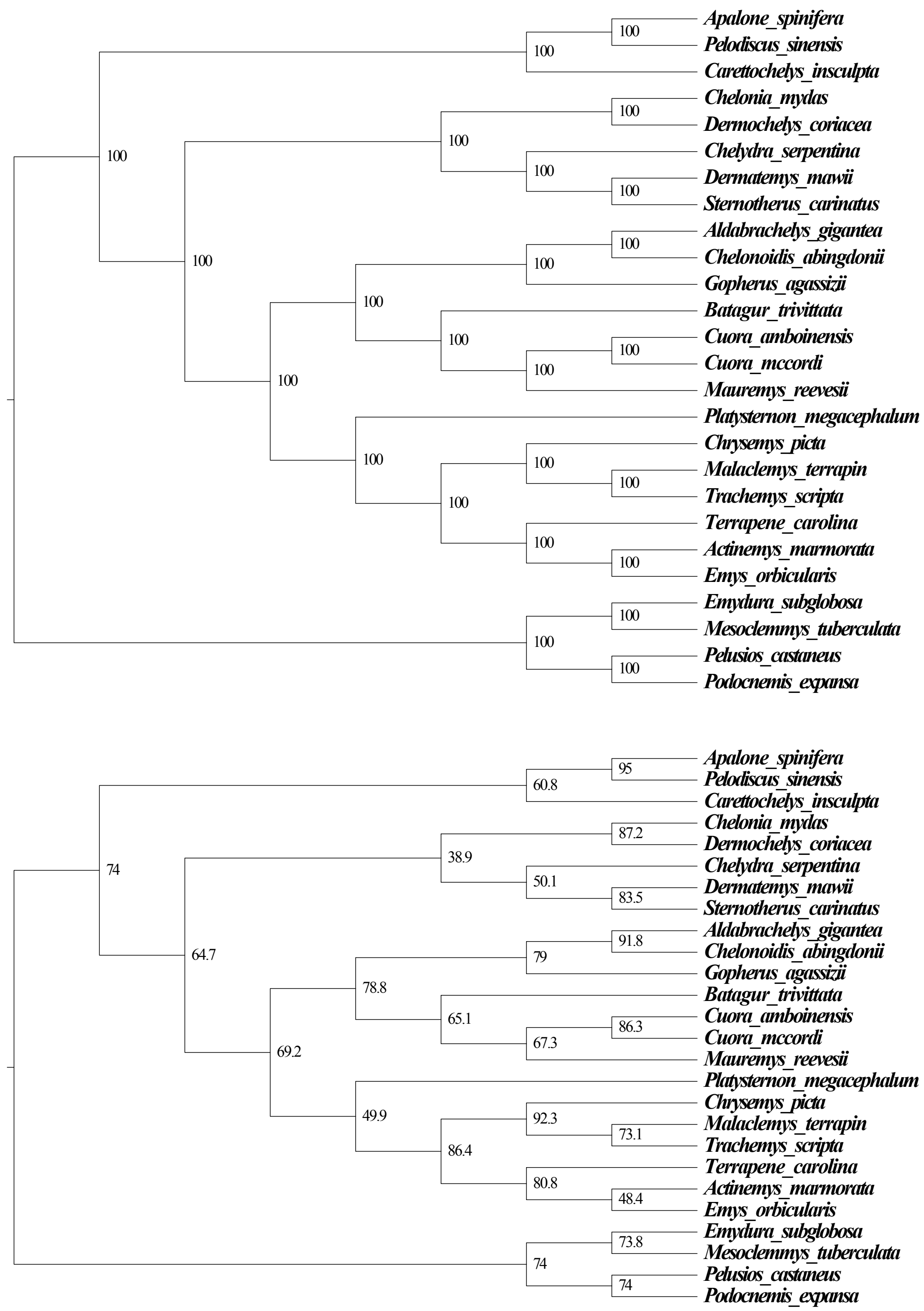

**Supplementary Figure 4.** SVDQuartets trees based on parsimony informative biallelic sites (CDS). Top: support values represent the proportion of 100 bootstrap replicates. Bottom: Site concordance factors for each node are given.

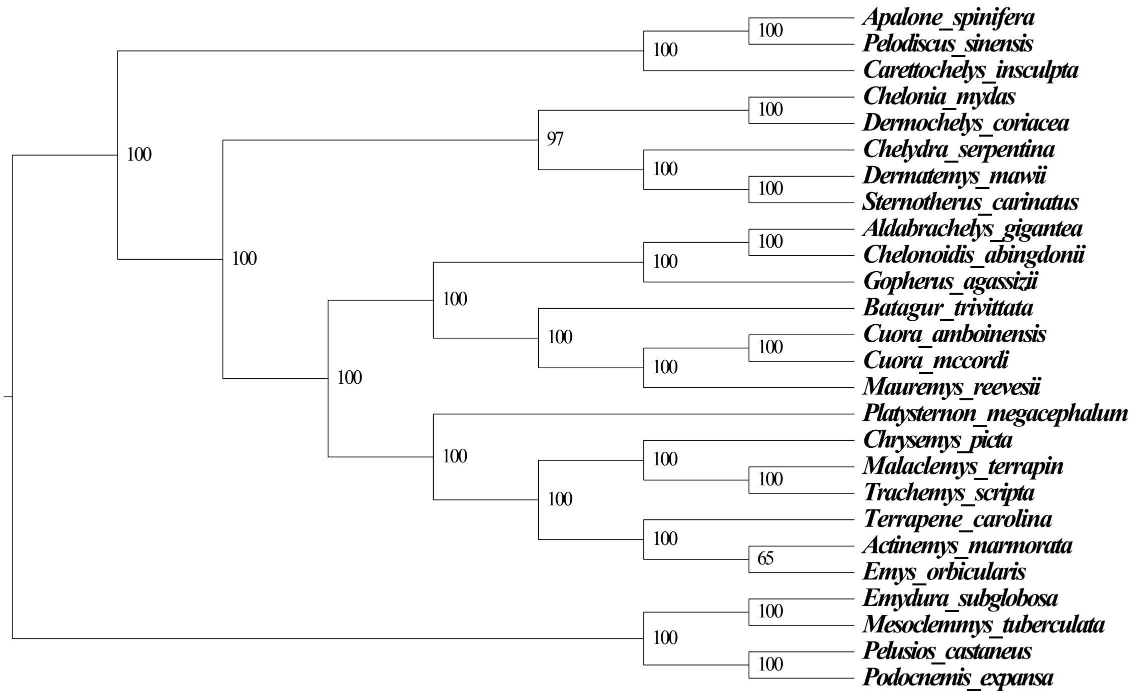


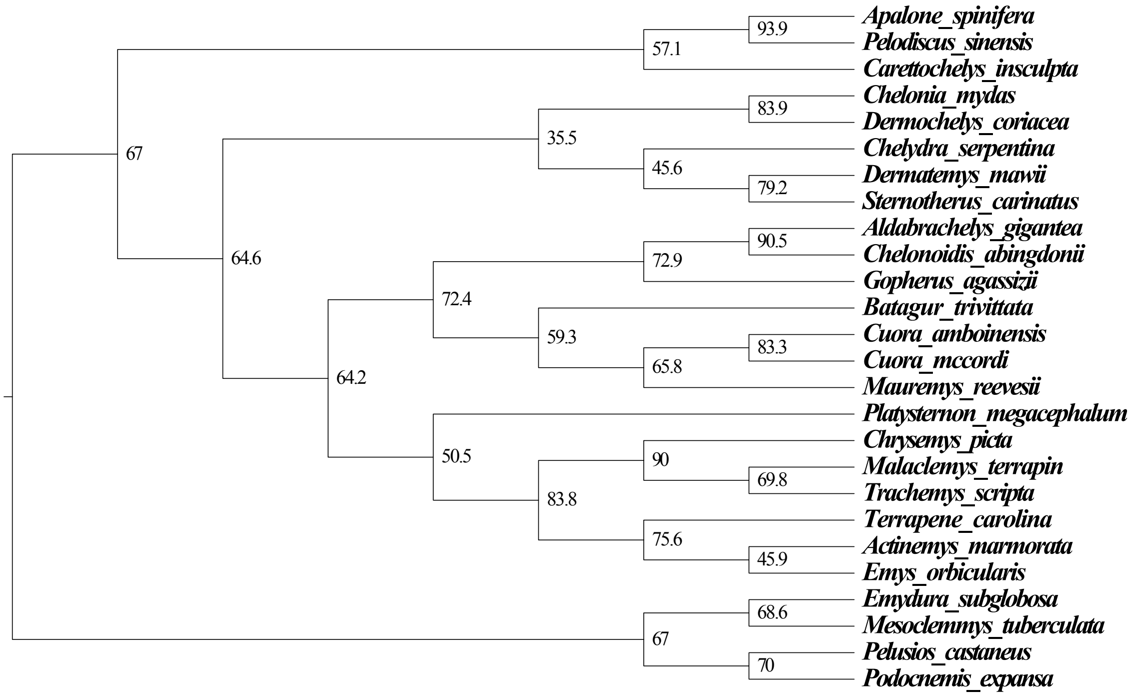

**Supplementary Figure 5.** SVDQuartets trees based on parsimony informative biallelic sites (introns). Top: support values represent the proportion of 100 bootstrap replicates. Bottom: Site concordance factors for each node are given.

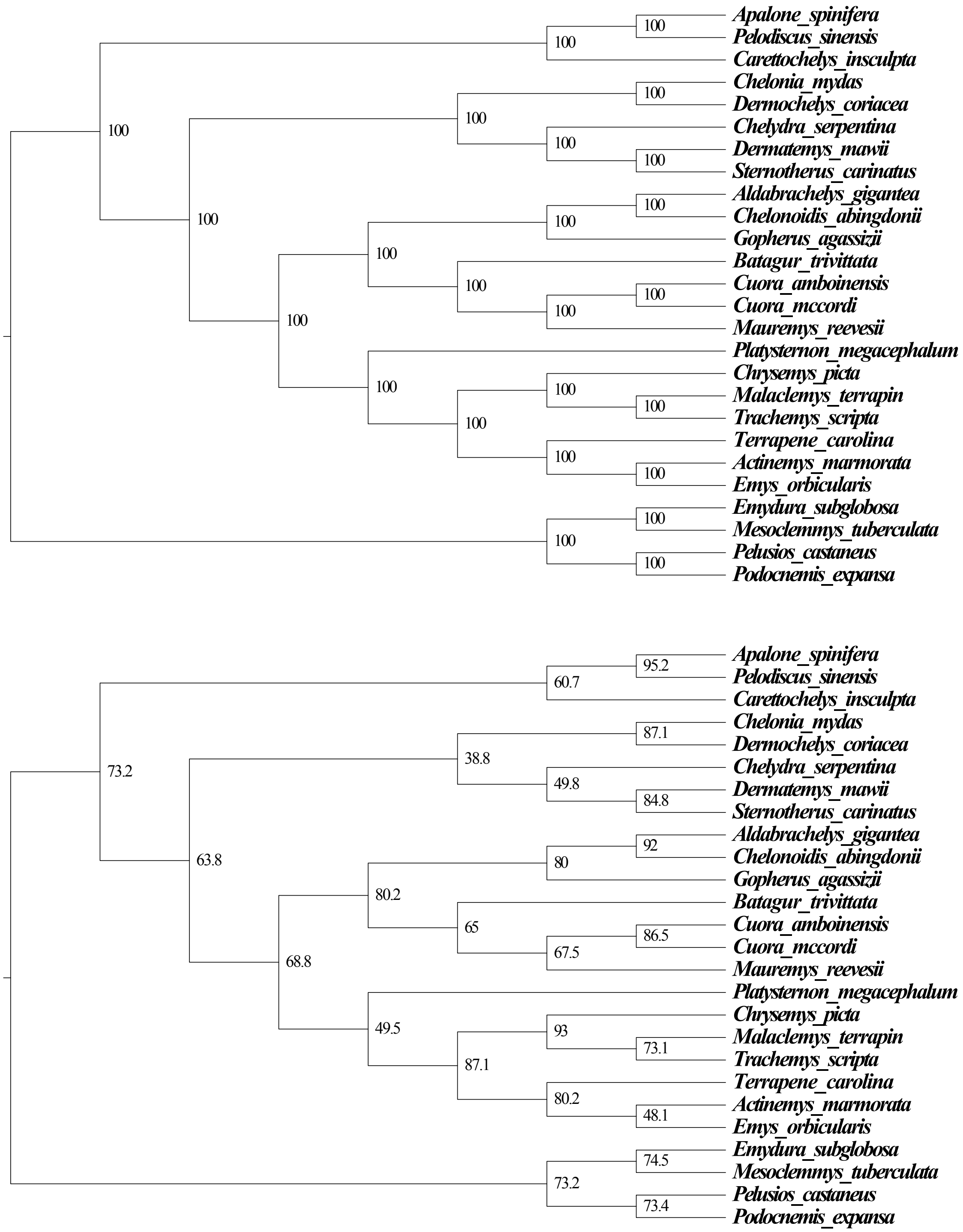

**Supplementary Figure 6.** SVDQuartets trees based on parsimony informative biallelic sites (5'-UTR). Top: support values represent the proportion of 100 bootstrap replicates. Bottom: Site concordance factors for each node are given.

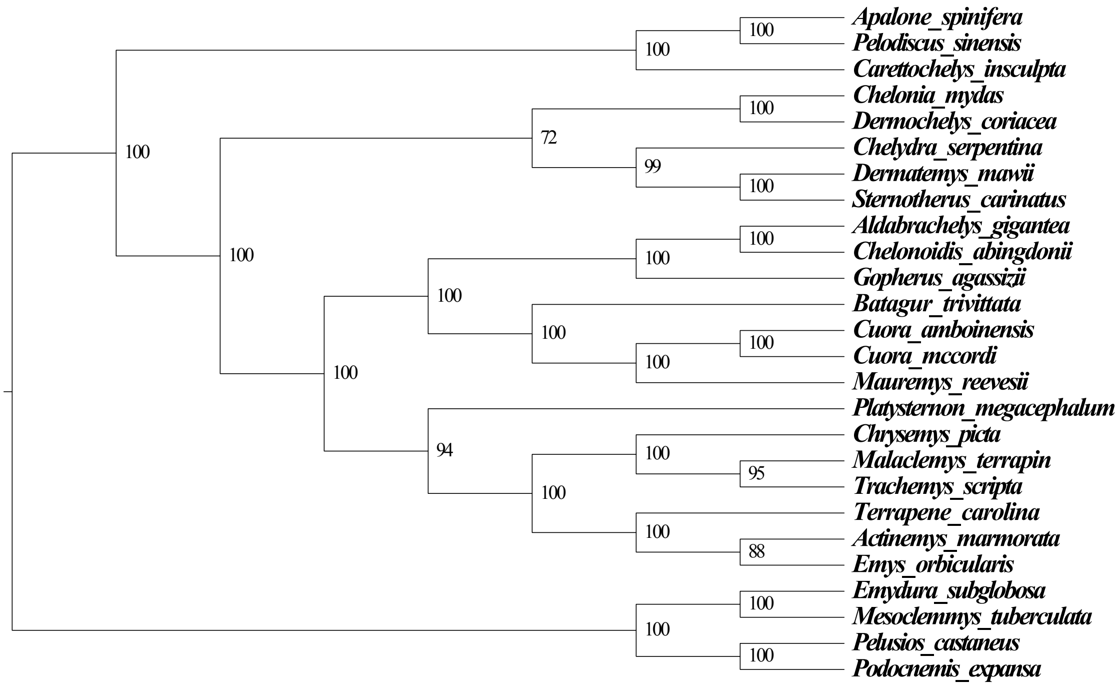


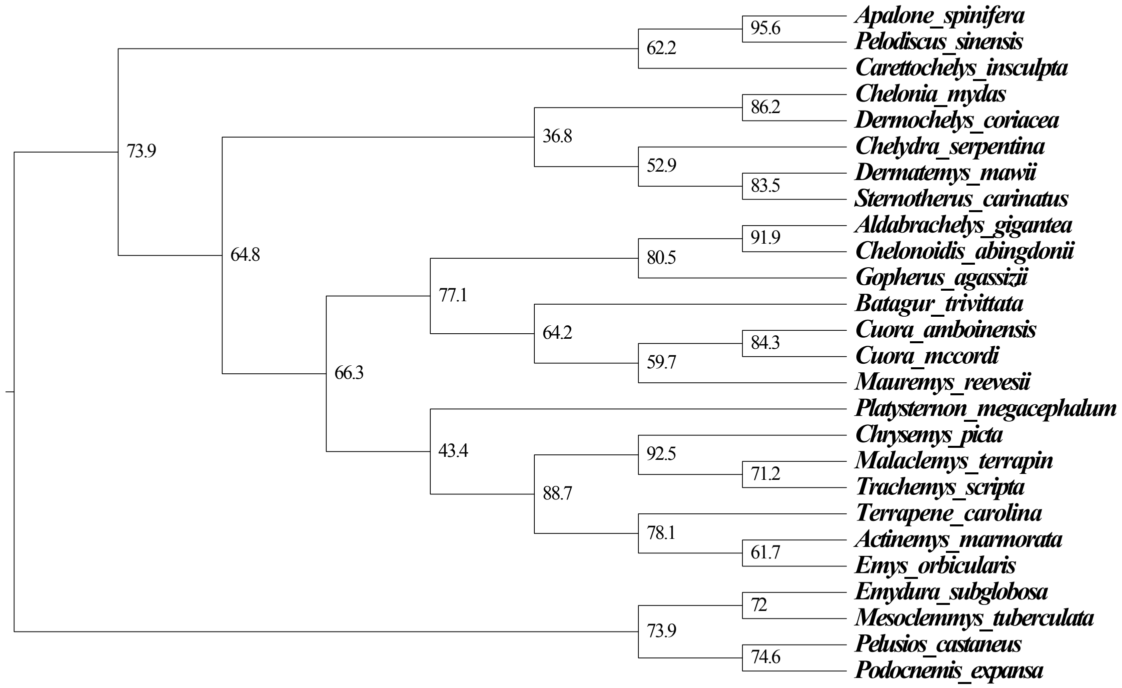

**Supplementary Figure 7.** SVDQuartets trees based on parsimony informative biallelic sites (3'-UTR). Top: support values represent the proportion of 100 bootstrap replicates. Bottom: Site concordance factors for each node are given.

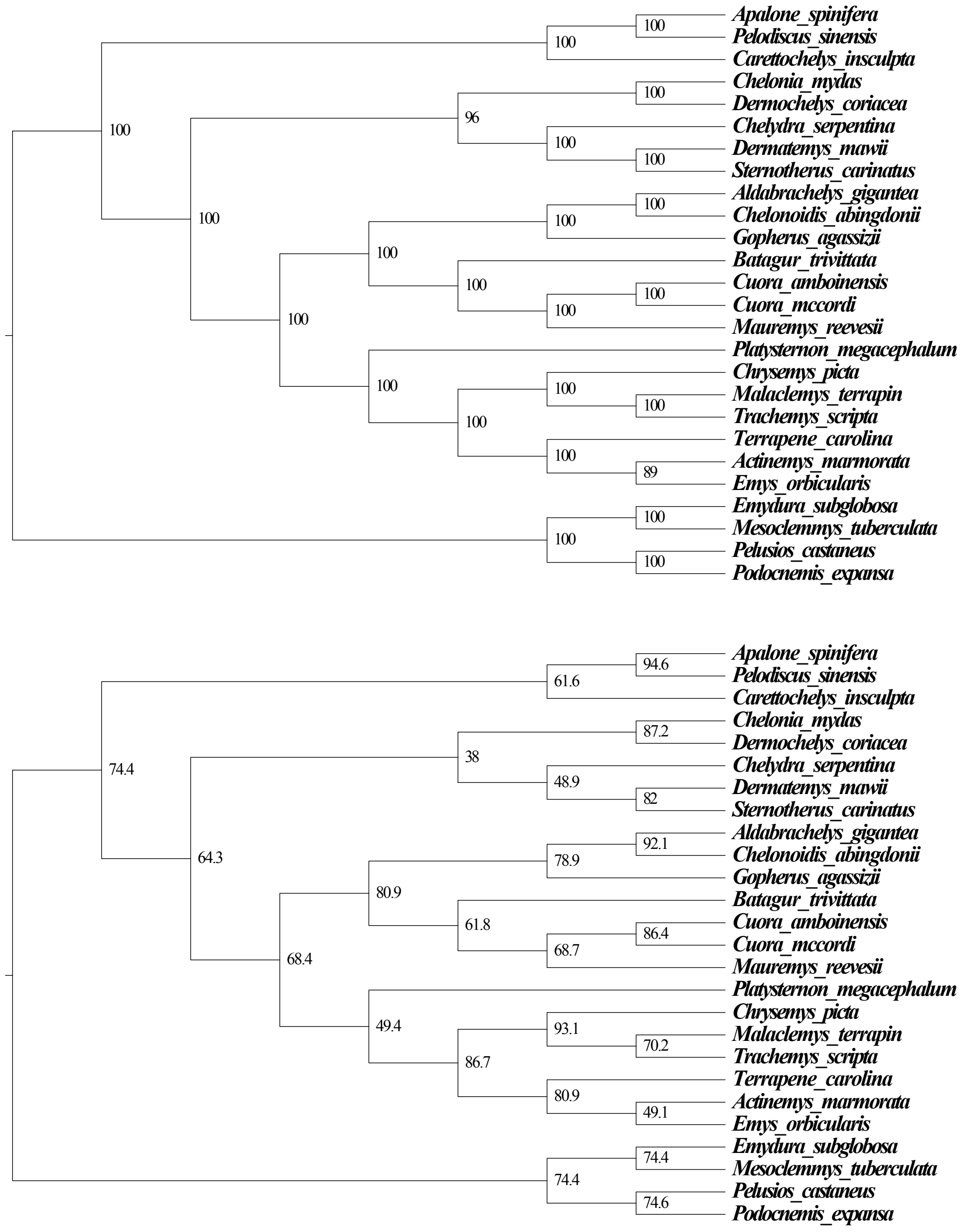

**Supplementary Figure 8.** SVDQuartets trees based on parsimony informative biallelic sites (genic_other). Top: support values represent the proportion of 100 bootstrap replicates. Bottom: Site concordance factors for each node are given.

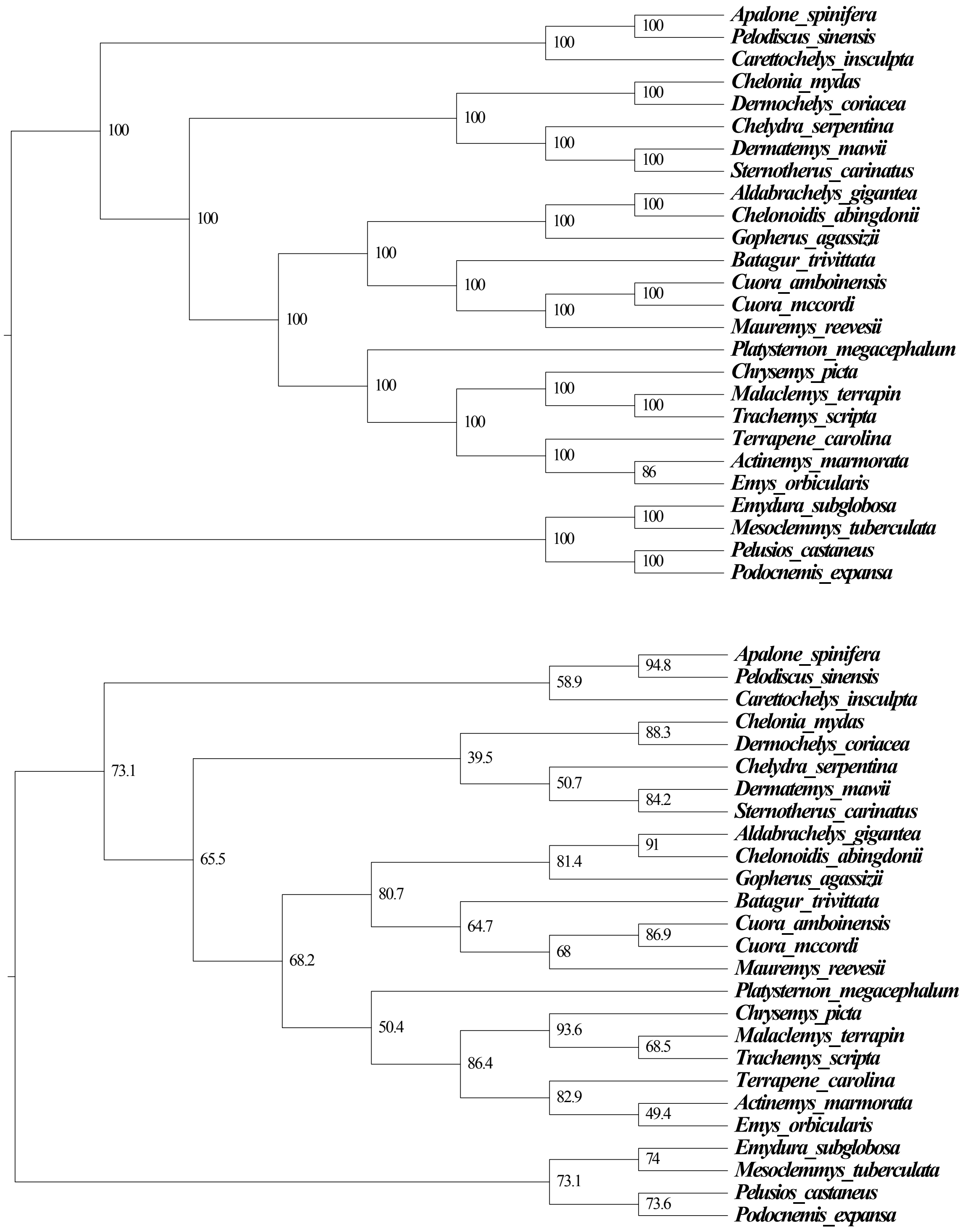

**Supplementary Figure 9.** SVDQuartets trees based on parsimony informative biallelic sites (lncRNA). Top: support values represent the proportion of 100 bootstrap replicates. Bottom: Site concordance factors for each node are given.

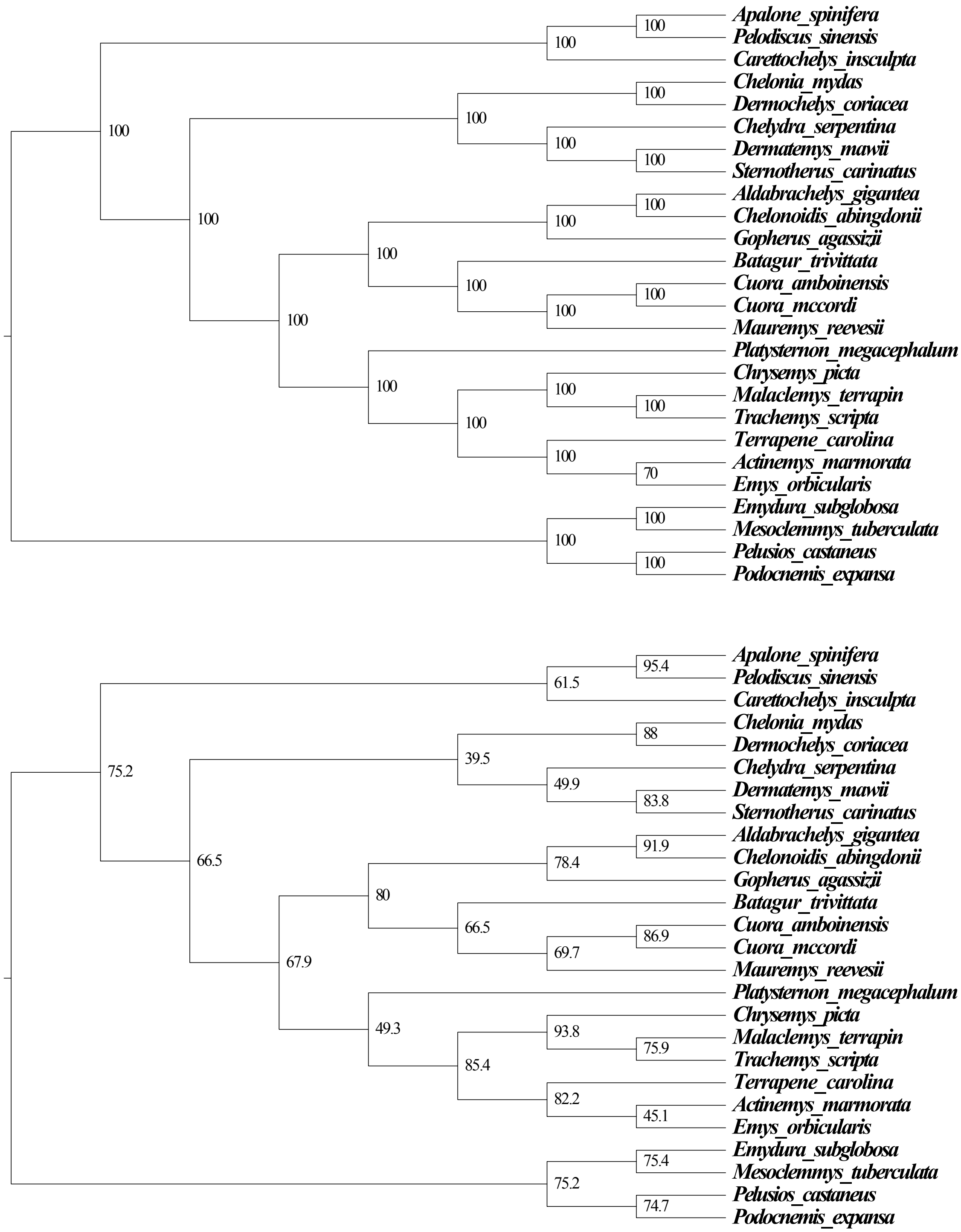

**Supplementary Figure 10.** SVDQuartets trees based on parsimony informative biallelic sites (pseudogenes). Top: support values represent the proportion of 100 bootstrap replicates. Bottom: Site concordance factors for each node are given.

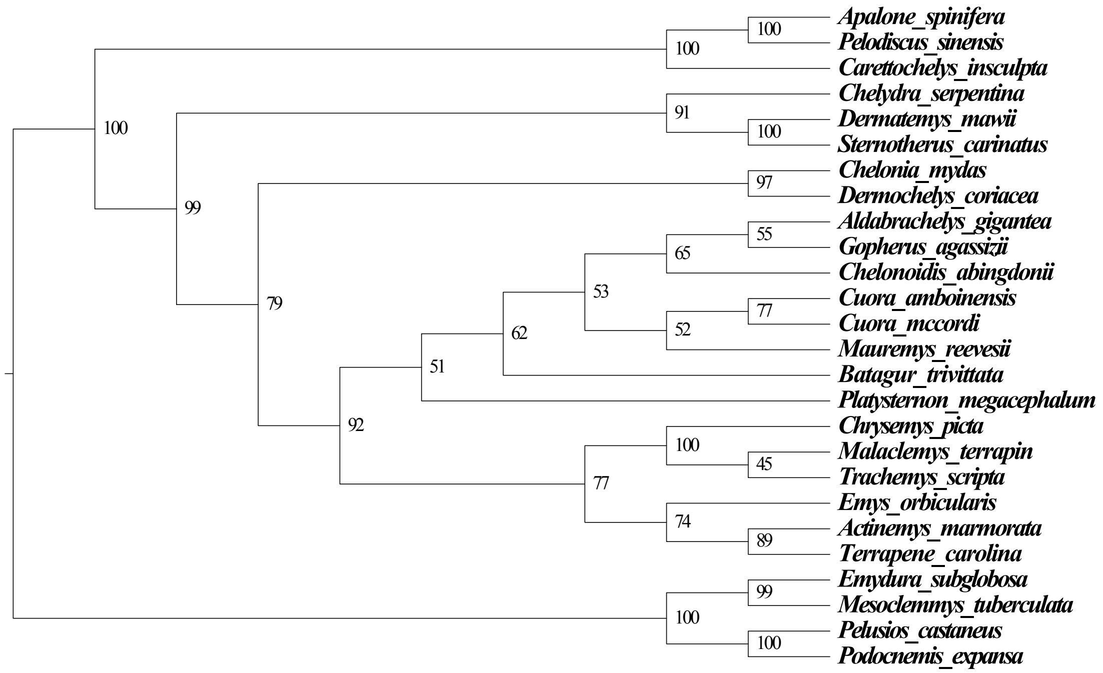


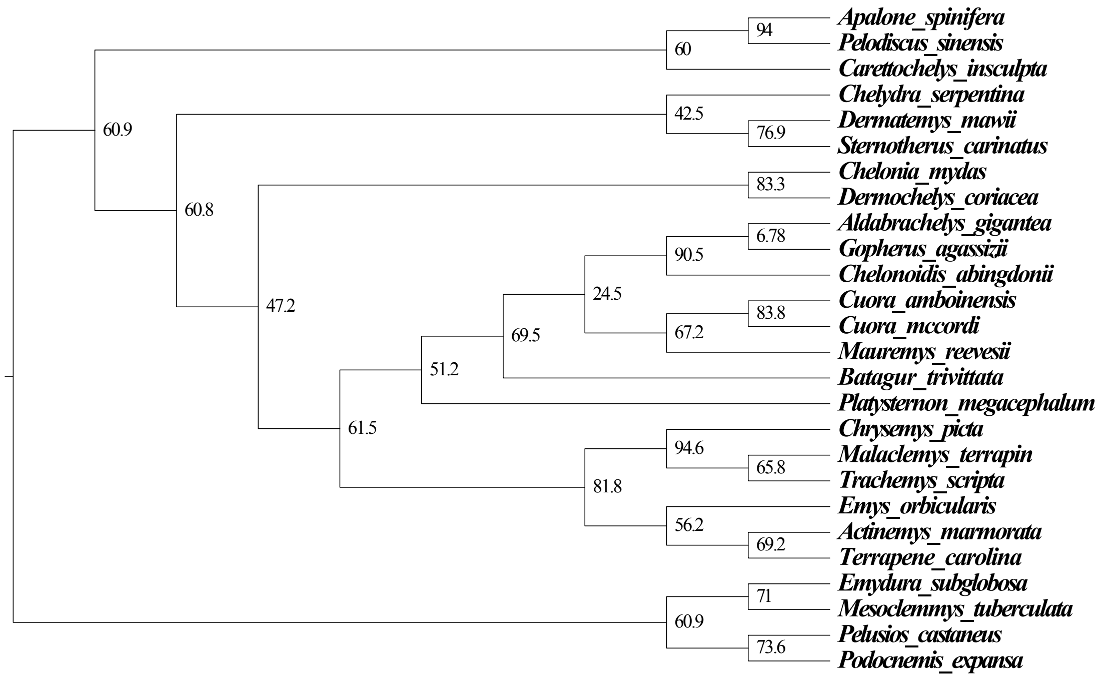

**Supplementary Figure 11.** SVDQuartets trees based on parsimony informative biallelic sites (smRNA). Support values represent the proportion of 100 bootstrap replicates. Site concordance factors could not be calculated due to polytomies.

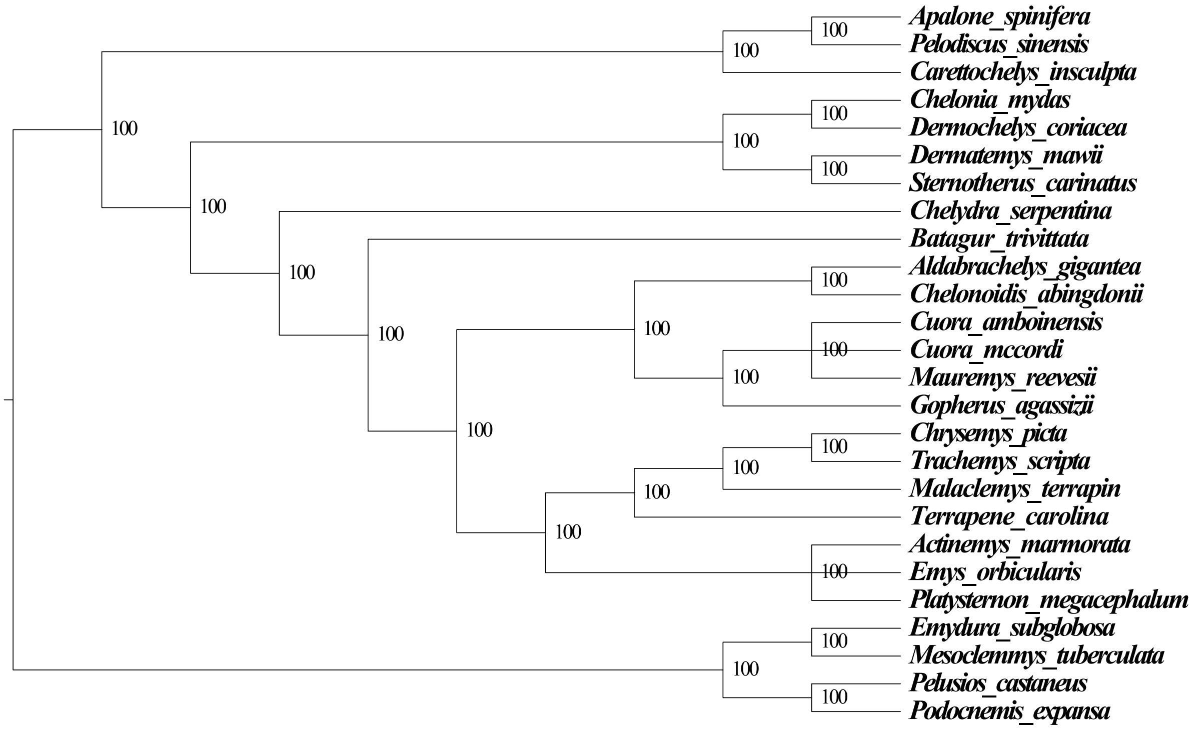

**Supplementary Figure 12.** Heat map of Robinson-Fould's distances between SVDQuartets trees inferred from parsimony informative biallelic sites from different locus types.

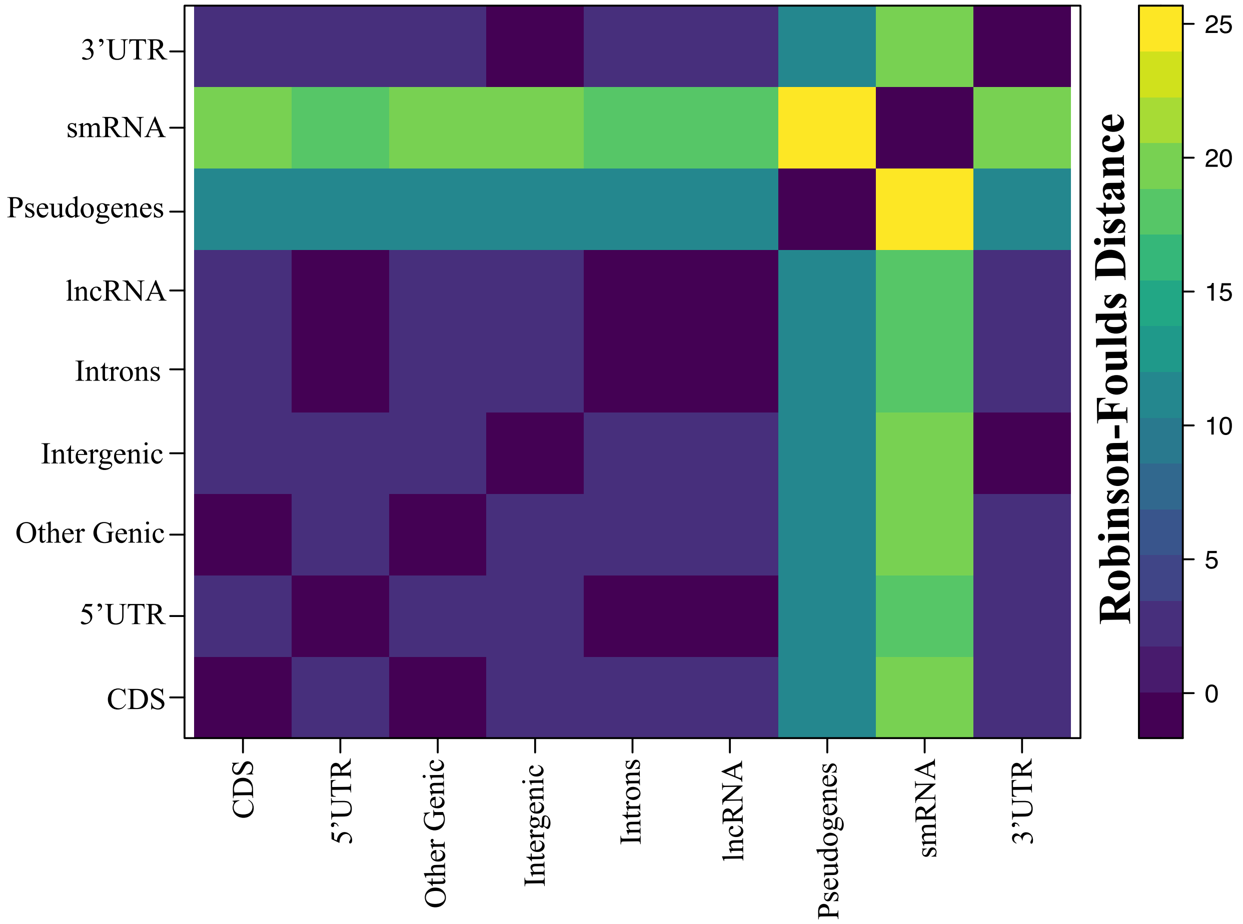

**Supplementary Figure 13.** Distribution of saturation coefficients for 685 single-copy ortholog alignments with complete taxon sampling.

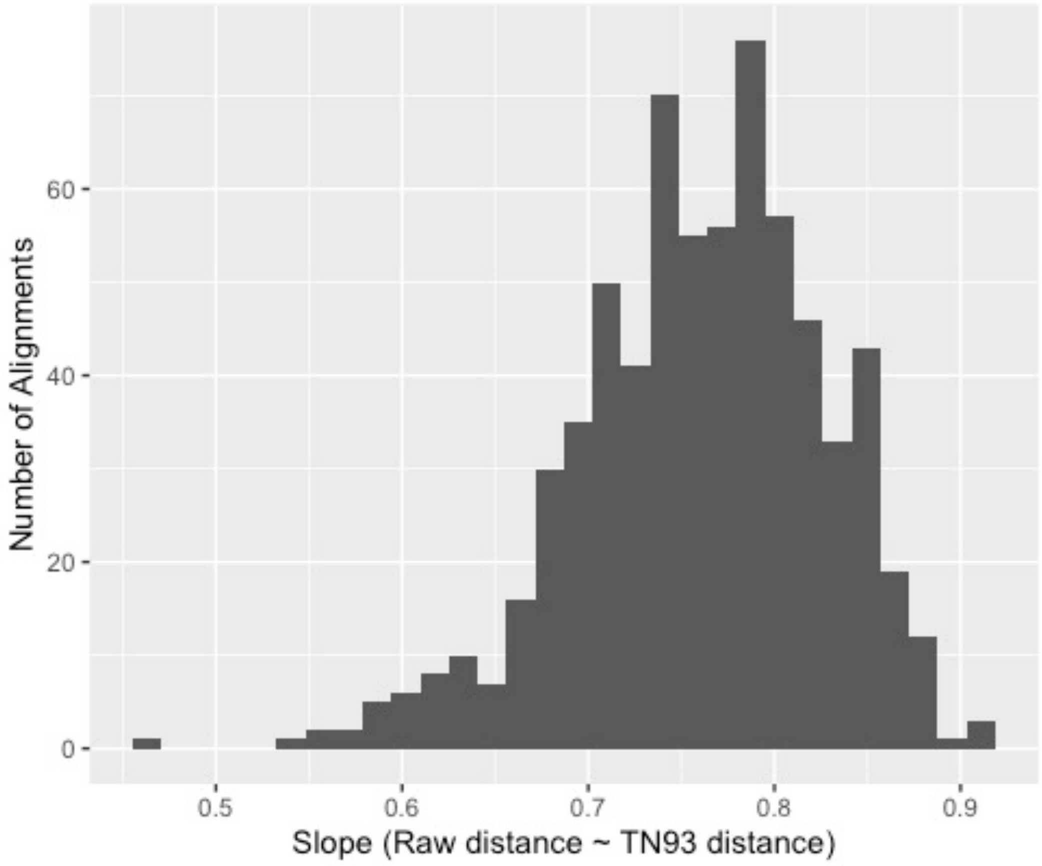

**Supplementary Figure 14.** Saturation plots by codon position in 685 single-copy ortholog alignments with complete taxon sampling.

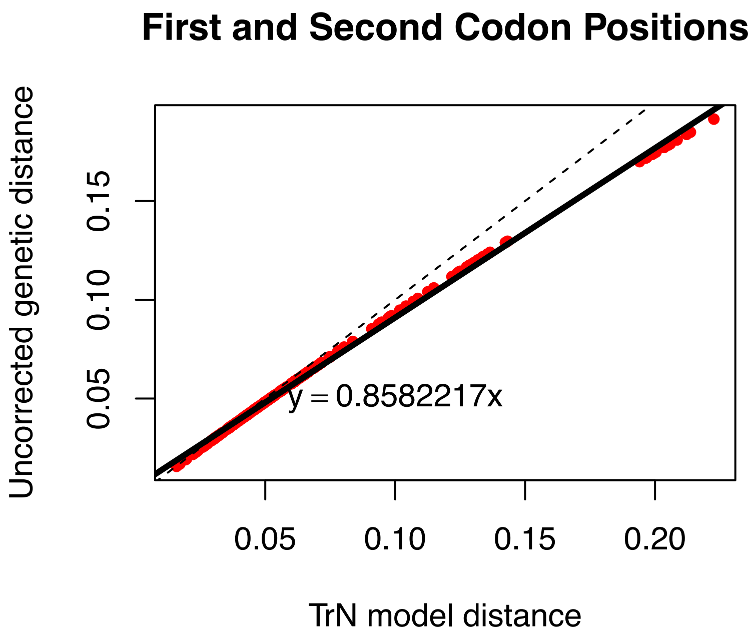


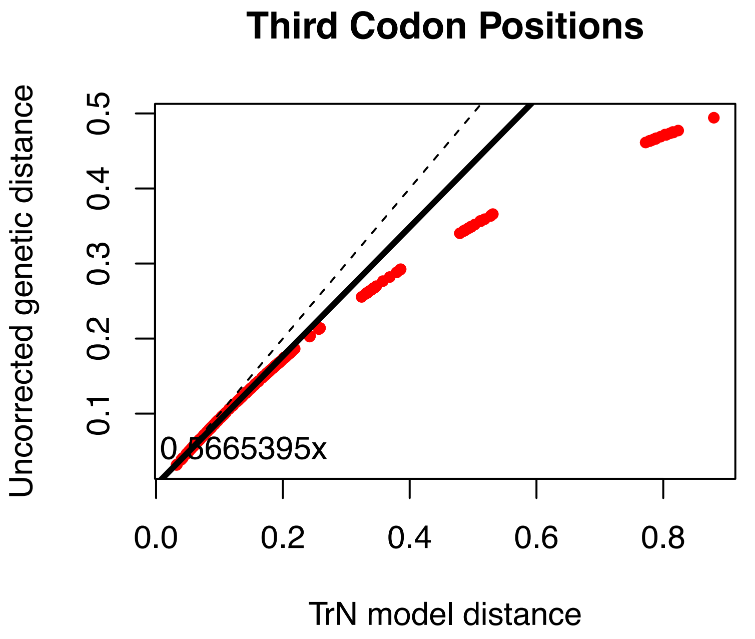

**Supplementary Figure 15.** Correspondence analyses results of Synonymous Codon Usage (RSCU) of the concatenated alignment of 685 single-copy orthologs with complete taxon sampling. Results by (a) species and (b) codons.

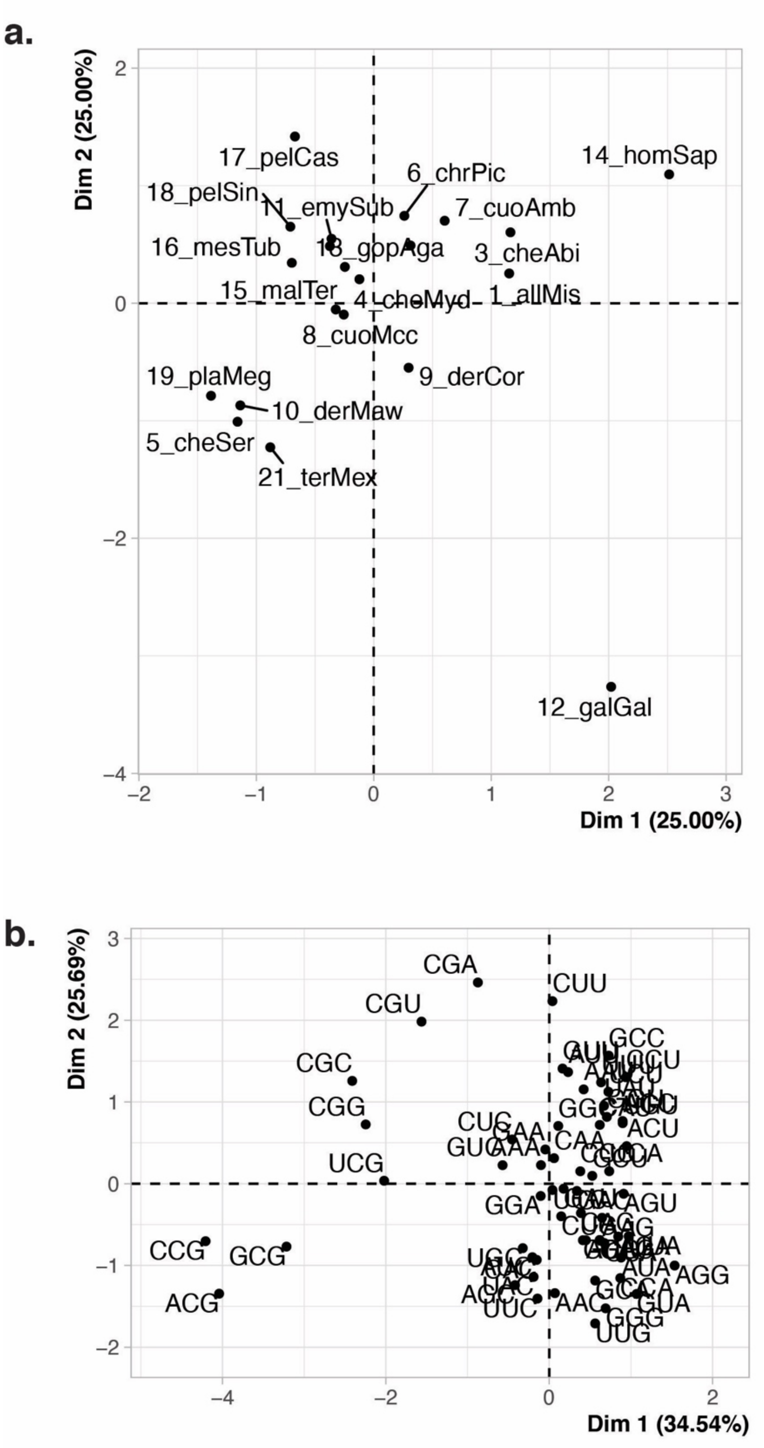

**Supplementary Figure 16.** Site concordance and estimated node ages in millions of years ago (M.Y.A.) from Thomson et al. (2021) in turtle evolution based on mapped SISRS data. Divergence times from Thomson et al. (2021) are in the Supplementary Data.

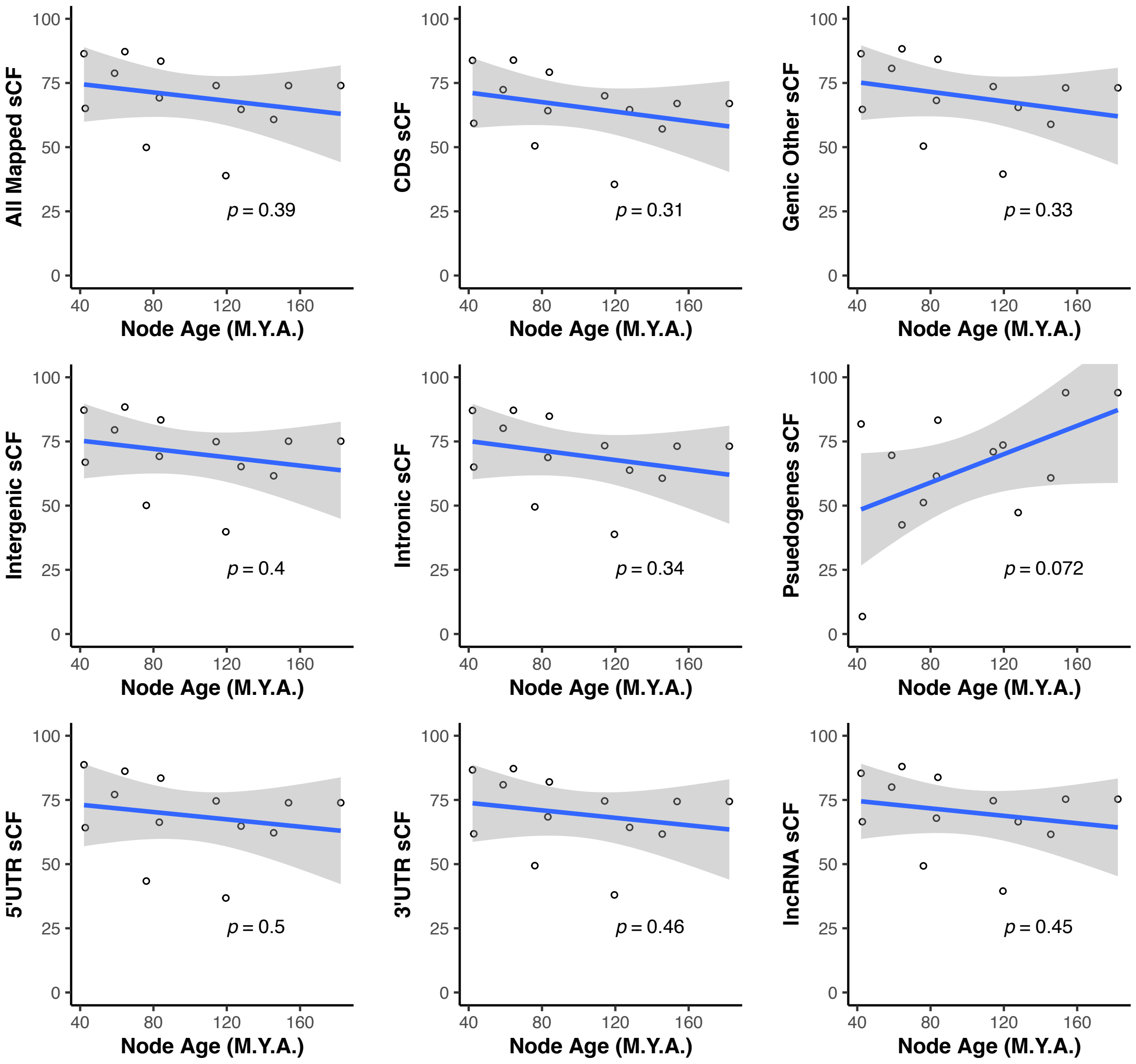

**Supplementary Figure 17.** Site concordance and estimated substitution rates in turtle evolution based on mapped SISRS data.

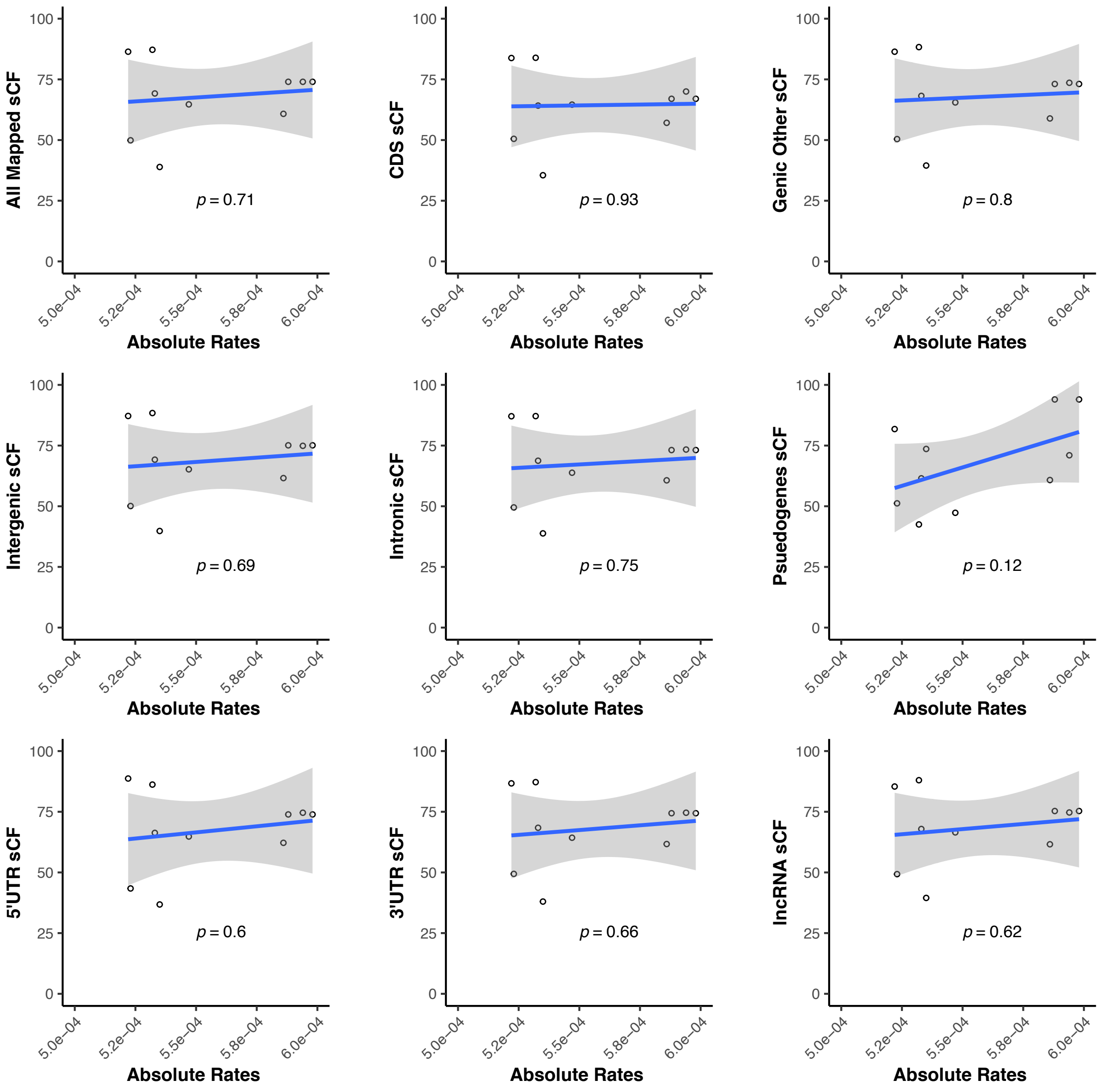

**Supplementary Figure 18.** Gene and site concordance (gCF and sCF), estimated node ages, and estimated substitution rates in turtle evolution based on single-copy orthologs and the concatenated partitioned maximum likelihood tree.

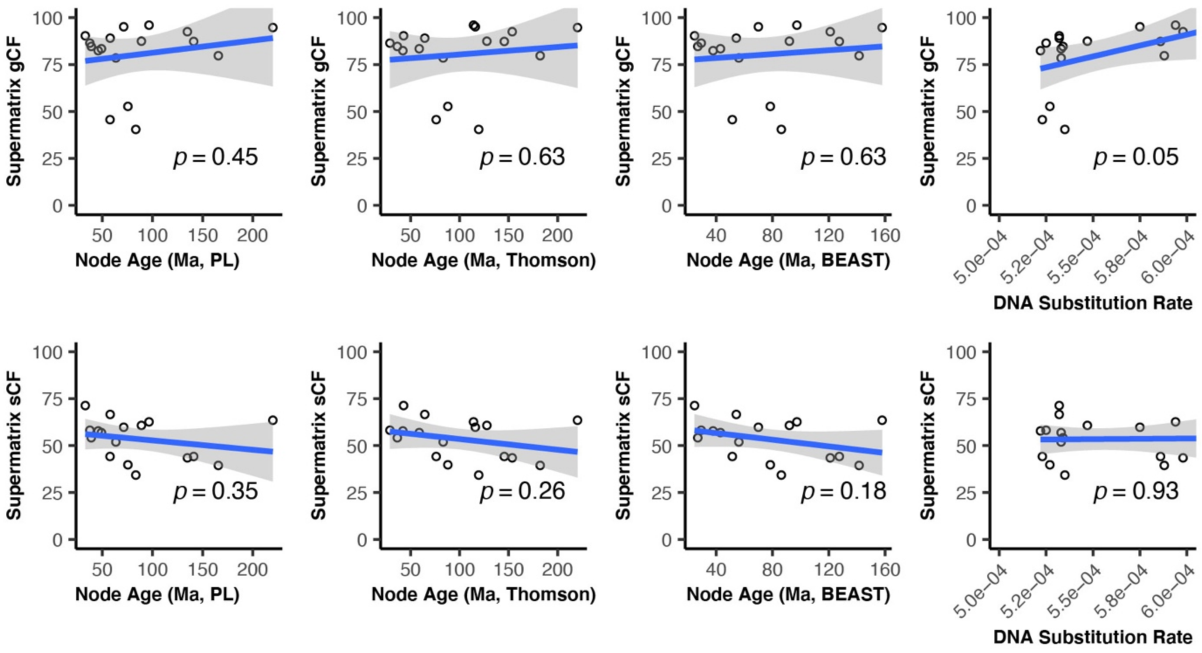
